# Cross-dataset reproducibility of human retinotopic maps

**DOI:** 10.1101/2021.04.12.439348

**Authors:** Marc M. Himmelberg, Jan W. Kurzawski, Noah C. Benson, Denis G. Pelli, Marisa Carrasco, Jonathan Winawer

## Abstract

Population receptive field (pRF) models fit to fMRI data are used to non-invasively measure retinotopic maps in human visual cortex, and these maps are a fundamental component of visual neuroscience experiments. Here, we examined the reproducibility of retinotopic maps across two datasets: a newly acquired retinotopy dataset from New York University (NYU) (n=44) and a public dataset from the Human Connectome Project (HCP) (n=181). Our goal was to assess the degree to which pRF properties are similar across datasets, despite substantial differences in their experimental protocols. The two datasets simultaneously differ in their stimulus apertures, participant pool, fMRI protocol, MRI field strength, and preprocessing pipeline. We assessed the cross-dataset reproducibility of the two datasets in terms of the similarity of vertex-wise pRF estimates and in terms of large-scale polar angle asymmetries in cortical magnification. Within V1, V2, V3, and hV4, the group-median NYU and HCP vertex-wise polar angle estimates were nearly identical. Both eccentricity and pRF size estimates were also strongly correlated between the two datasets, but with a slope different from 1; the eccentricity and pRF size estimates were systematically greater in the NYU data. Next, to compare large-scale map properties, we quantified two polar angle asymmetries in V1 cortical magnification previously identified in the HCP data. The NYU dataset confirms earlier reports that more cortical surface area represents horizontal than vertical visual field meridian, and lower than upper vertical visual field meridian. Together, our findings show that the retinotopic properties of V1, V2, V3, and hV4 can be reliably measured across two datasets, despite numerous differences in their experimental design. fMRI-derived retinotopic maps are reproducible because they rely on an explicit computational model of the fMRI response. In the case of pRF mapping, the model is grounded in physiological evidence of how visual receptive fields are organized, allowing one to quantitatively characterize the BOLD signal in terms of stimulus properties (i.e., location and size). The new NYU Retinotopy Dataset will serve as a useful benchmark for testing hypotheses about the organization of visual areas and for comparison to the HCP 7T Retinotopy Dataset.

## 1. Introduction

The human visual system preserves image structure in cortical maps of the visual field. These retinotopic maps are the basis by which measurements across labs, populations, and tasks are compared. Functional magnetic resonance imaging (fMRI) has made it possible to non-invasively measure retinotopic maps from visually evoked activity in human cortex (DeYoe et al., 1994; Engel et al., 1994; Sereno et al., 1995). Methodological advances have led to sophisticated computational models such as population receptive field (pRF) modeling (Dumoulin & Wandell, 2008), which has become a core component of experiments investigating human visual cortex (reviewed by Wandell & Winawer, 2015).

Recently, the Human Connectome Project (HCP) 7 Tesla Retinotopy Dataset has been made available for public use (Benson et al., 2018; Van Essen et al., 2012). This dataset contains high-quality retinotopy data from 181 participants, collected with 7T magnetic resonance imaging (MRI), with 30 minutes of data per participant. The high quality and large size of the dataset make it an attractive resource for researchers to address a wide variety of questions. It has been used to investigate retinotopic organization in visual cortex, including the organization of core pRF parameters (Benson et al., 2018), the organization of white matter connections between thalamus and foveal and peripheral representations of extrastriate cortex (Kurzawski et al., 2020), the identification new retinotopic maps in the cerebellum (van Es et al., 2019), hippocampus (Silson et al., 2021), the visual organization in the default network (Szinte & Knapen, 2020), and polar angle meridian asymmetries in cortical magnification in primary visual cortex (V1) (Benson et al., 2021). One benefit of the HCP dataset is that its large size permits researchers to conduct checks of internal reliability. For example, if a finding is observed in one half of the participant sample, is it also observed in the other half? However, these internal checks rely on data that have been collected using the same MRI scanner, preprocessed via the same pipeline, and analyzed using the same pRF software. Therefore, systematic biases affecting a full dataset could make the data internally reliable, but not generalizable.

Visual neuroscientists assume that retinotopic organization is broadly consistent across the healthy human population. For example, there is no uncertainty about V1’s localization in the calcarine sulcus or that receptive field sizes increase with eccentricity. Previous studies have quantified the reliability of retinotopic properties within their own datasets (Benson et al., 2018; Binda et al., 2013; Senden et al., 2014; van Dijk et al., 2016; Zeidman et al., 2018; Zuiderbaan et al., 2012). However, it is important to validate scientific findings relating to retinotopic organization by testing the extent to which they can be reproduced in *independent* datasets, especially in an age of open science and data-sharing, where visual neuroscientists are granted access to an increasing number of large, public datasets that differ in many ways. One robust assessment of the validity of such findings is via their reproduction in a retinotopy dataset that uses a different MRI scanner, fMRI protocol, stimulus aperture, and preprocessing pipeline. Such differences are typical in neuroimaging studies across labs. An explicit test of the reproducibility of retinotopic maps between large datasets that simultaneously differ in many aspects has not been conducted. To measure the reliability of retinotopic maps across independent datasets, we have acquired, and made publicly available, a large new set of retinotopic data — the New York University (NYU) Retinotopy Dataset — that differs from the HCP 7T Retinotopy Dataset in many ways.

This paper has three goals. The first is to describe and provide a public release of a high-quality retinotopy dataset that can be used by other researchers to address open questions about the organization of retinotopic maps in the human brain. The second is to quantify the cross-dataset reproducibility of pRF estimates at the vertex scale, after cross-subject alignment of cortical surfaces. The third is to quantify the cross-dataset reproducibility of polar angle asymmetries in V1 surface area.

First, we present visualizations and a description of the new NYU Retinotopy Dataset at the individual and group level. Retinotopic maps are differentiated based on polar angle and eccentricity representations of the visual field (Dumoulin & Wandell, 2008; Wandell et al., 2007). Thus, we second consider the cross-dataset reproducibility of vertex-wise pRF estimates (polar angle, eccentricity, and pRF size) between the NYU and HCP data. We do so in four visual areas: V1, V2, V3, and hV4. To make vertex-wise comparisons between the two datasets, all individual cortical surfaces are aligned to a standardized, common template surface so that the datasets can be compared at the mm scale, and the native geometry of the individuals is discarded. We then compare pRF parameters at each vertex between the two datasets. Third, we examine large scale properties of the maps in terms of cortical surface area. Findings from a subset of data from the full HCP dataset (n=163 of 181) show large-scale polar angle asymmetries in V1 cortical magnification: substantially more cortical surface area is dedicated to processing the horizontal than the vertical visual field meridian (the horizontal-vertical anisotropy; HVA), and more cortical surface area is dedicated to the lower than the upper vertical visual field meridian (the vertical meridian asymmetry; VMA) (Benson et al., 2021). These cortical magnification asymmetries are derived from both pRF estimates and surface area measurements. Here, we assess the HVA and VMA in V1 cortical surface area in our new (NYU) dataset. The cortical magnification measures reflect the relation between retinotopic coordinates and cortical geometry (local surface area) and therefore, unlike the vertex-wise comparisons, require analysis on each participant’s native cortical surface.

## 2. Methods

### 2.1 Participants

Forty-four participants (25 females, 19 males, mean age=28.8 years) were recruited from New York University (NYU). All participants had normal or corrected-to-normal vision and completed a 1 – 1.5-hour scanning session. All participants provided written informed consent and approved the public release of anonymized data. The experiment was conducted in accordance with the Declaration of Helsinki and was approved by the NYU ethics committee on activities involving human participants.

### 2.2 fMRI stimulus display

Participants viewed the pRF stimulus in the MRI scanner using a ProPixx DLP LED projector (VPixx Technologies Inc., Saint-Bruno-de-Montarville, QC, Canada). The stimulus image was projected onto an acrylic back-projection screen (60 cm x 36.2 cm) in the scanner bore. The projected image had a resolution of 1920 × 1080 and a refresh rate of 60 Hz. The display was calibrated using a linearized lookup table and the display luminance was 500 cd/m^2^. Participants viewed the screen at a distance of 83.5 cm (from eyes to the screen) using an angled mirror that was mounted on the head coil.

### 2.3 PRF Stimulus

The pRF stimulus was generated in MATLAB 2017a and was presented using the Psychophysics Toolbox v3 (Kleiner et al., 2007) and custom *vistadisp* software (https://github.com/WinawerLab/vistadisp) on an iMac computer. Stimulus image patterns were shown within a bar aperture that swept across the screen throughout each scan. There were 100 stimulus image patterns. The stimulus image patterns were identical to those used in the HCP 7T Retinotopy Dataset, except for rescaling to the size of our display. Each image pattern was composed of colorful objects, faces, and scenes at multiple scales (Kriegeskorte, 2008) that were superimposed on an achromatic pink-noise (1/f) background (see **Figure 1A** for example of the pRF stimulus). The stimulus image pattern was windowed within a circular aperture (12.4° radius) and was revealed through the bar aperture that swept across the screen in 24 equal steps, once per second, synchronized to the MR image acquisition (TR 1 second). The bar aperture was superimposed on a polar fixation grid placed upon a uniform gray background, with a red or green dot at the center (3 pixels, or 0.07°). The polar grid and small fixation point were used to encourage good fixation behavior, as previously described (Schira et al., 2009). The bar aperture was 3.1° in width, which was ⅛-th of the full stimulus extent (24.8° diameter). The bar aperture swept across the screen in 8 directions (see **Figure 1B** for sweep direction order and timings). Each sweep began at the edge of the circular aperture. Horizontal and vertical sweeps traversed the entire diameter of the aperture. Diagonal sweeps only traversed half of the aperture (the second half of the sweep was replaced with a blank gray screen, other than the fixation dot and grid). The blank periods help discriminate between nonvisual responses and responses from populations of neurons with exceptionally large receptive field sizes (Dumoulin & Wandell, 2008).

**Figure 1.**
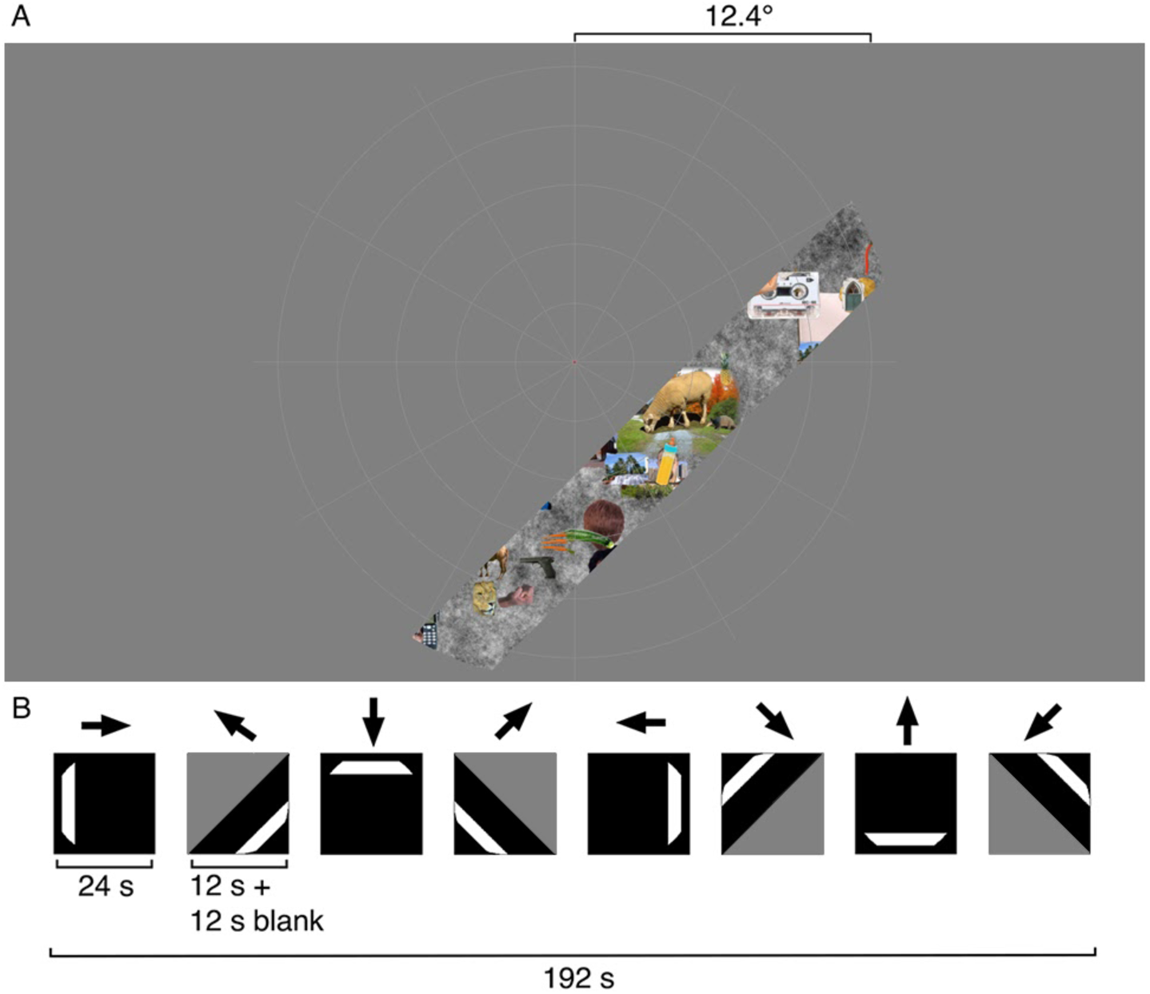
Example of the pRF stimulus and experimental design. (**A**) The pRF stimulus at a single time point. The bar aperture reveals the stimulus image pattern composed of colorful objects, faces, and scenes at different scales, randomly placed across an achromatic pink noise background. The fixation point at the center of the fixation grid changes from green to red and red to green at random times. (**B**) Experimental design for a single scan across 192 s – the pRF bar is swept across the screen in the eight cardinal directions, with each directional sweep lasting 24 s. Gray triangles indicate blank periods. Black regions correspond to the visual field covered by the sweeping bar (excluding the stimulus aperture).

Each directional sweep lasted 24 s, with the full stimulus run lasting 192 s. The stimulus image updated 3 times per second without intermediate blanks, thus the three images were shown for 0.333 s at each aperture position. Participants completed a fixation task to ensure that they were maintaining central fixation and remained alert throughout the scan. Participants were required to respond, via button press, when the central fixation dot changed from green to red, or vice versa. The full stimulus sequence was completed once per scan, and 4 to 12 scans were collected per participant, with the identical aperture sequence in each of the repeated scans.

### 2.4 Structural and functional data acquisition

Structural and functional data were acquired on a 3T Siemens MAGNETOM Prisma MRI scanner (Siemens Medical Solutions, Erlangen, Germany) at the Center for Brain Imaging at NYU. Both the structural and functional images were acquired using a Siemens 64-channel head coil. Between 1 and 2 full brain T1-weighted (T1w) MPRAGE anatomical images were acquired for each participant (TR, 2400 ms; TE, 2.4 ms; voxel size, 0.8mm^3^ isotropic; flip angle, 8°), and were auto-aligned to a template to ensure a similar slice prescription for all participants.

For a subset of participants (n=11) for whom there was sufficient time in the scanning session, a full brain T2-weighted (T2w) anatomical image was obtained to aid in automatic segmentation (TR, 3200 ms; TE, 564 ms; voxel size, 0.9mm^3^ isotropic; flip angle, 120°). Between 4 and 12 functional echo-planar images (EPIs) were acquired for each participant using a T2*-weighted multiband EPI sequence (TR, 1000 ms; TE, 37 ms; voxel size, 2mm^3^; flip angle, 68°; multiband acceleration factor, 6; phase-encoding, posterior-anterior) (Feinberg et al., 2010; Xu et al., 2013). Additionally, two distortion maps were acquired to correct susceptibility distortions in the functional images: one spin-echo image with anterior-posterior (AP) and one with posterior-anterior (PA) phase encoding.

### 2.5 Preprocessing of structural and functional data using fMRIPrep

Anatomical and functional preprocessing was performed using *fMRIPrep* v.20.0.1 (Esteban et al., 2019; Gorgolewski et al., 2011). Each T1w anatomical image was corrected for intensity inhomogeneity and was skull-stripped. Automatic brain segmentation of cerebrospinal fluid, white-matter, and gray-matter was performed on the skull-stripped T1w image using *fast* (Zhang et al., 2001). If acquired, the T2w image was included as an additional input for brain segmentation. Cortical surfaces were then reconstructed using Freesurfer’s *recon-all* (Dale et al., 1999) and an estimated brain mask was refined using a custom variation of the method.

For each participant’s functional images, the following preprocessing was performed. A reference volume and its skull-stripped version were created using the custom methodology from *fMRIPrep*. A B0-nonuniformity map was estimated based on the two spin-echo images with opposing phase-encoding directions (i.e., the AP and PA distortion maps). Using the estimated distortion, a corrected functional reference image was calculated to ensure accurate co-registration with the anatomical reference. The functional reference was co-registered to the T1w anatomical reference. Co-registration was configured with six degrees of freedom.

Head-motion parameters with respect to the functional reference (transformation matrices) were estimated first. The functional images were then slice-time corrected in which all slices were realigned in time to the middle of each TR. The resampling of the slice-time corrected functional data to the T1w anatomical space was performed in a one-shot interpolation by composing all the pertinent transformations (i.e., head-motion transform matrices, susceptibility distortion correction, and coregistration to the T1w anatomical space). The single interpolation step reduces the effect of blurring the signal multiple times across multiple transformations.

Finally, these pre-processed time-series data were resampled onto the *fsnative* surface using Freesurfer’s *mri_surf2vol* by averaging across the cortical ribbon.

### 2.6 Population receptive field model

PRF analysis was conducted using *vistasoft* (https://vistalab.stanford.edu/software/, Vista Lab, Stanford University). Here, a pRF is modelled as a circular 2D-Gaussian, as described in Dumoulin and Wandell (2008). The Gaussian is parameterized by values at each vertex for *x*, *y*, and *σ*. The *x* and *y* parameters specify the center position of the 2D-Gaussian in the visual field. The *σ* parameter, the standard deviation of the 2D-Gaussian, specifies the size (or spread) of the receptive field. The 2D-Gaussian is multiplied pointwise by the stimulus contrast aperture, and then convolved with a hemodynamic response function (HRF) to predict the BOLD percent signal change (or ‘BOLD signal’). The HRF is parameterized by 5 values, describing a difference of two gamma functions, as used previously (Dumoulin & Wandell, 2008; Friston et al., 1998; Harvey & Dumoulin, 2011; Worsley et al., 2002). The HRF was assumed to be the same across vertices within a participant but differed among participants.

The *vistasoft* pRF model was implemented using the *prf-analyze-vista* docker container (https://github.com/vistalab/prfmodel; (Lerma-Usabiaga et al., 2020). The software finds the optimal pRF parameters for each vertex, and the optimal HRF parameters averaged across vertices, by minimizing the residual sum of squares between the predicted time-series and the BOLD signal. This is completed using a multi-stage coarse-to-fine approach. The basic two-stage coarse-to-fine component is described in detail by Dumoulin and Wandell (2008), and the addition of the HRF fit is described in detail by Harvey and Dumoulin (2011). In brief, in the first stage, the data were temporally decimated by a factor of two to remove high frequency noise and the pRF parameters (*x*, *y*, and *σ*) were fit using a brute force grid search. The results of this fit were taken as the starting point of a second-stage search fit for these parameters using the full time-series. The pRF parameters were then held fixed and the HRF parameters were fit by a search, choosing the parameters that minimize the squared error between data and prediction averaged across vertices. Finally, the HRF parameters were held fixed and the pRF parameters were refit to the data.

We implemented the pRF model on three formats of data. First, we solved the pRF model for each vertex on individual participant data to produce pRF maps on *fsnative* surfaces. Second, for each participant, we interpolated the pRF maps by nearest neighbor from the *fsnative* space to the *fsaverage* space. This interpolation puts the pRF model solutions for all participants in the same brain space, enabling us to derive the vertex-wise median parameters across all participants. Third, we interpolated each participant’s average time-series from the *fsnative* to the *fsaverage* surface, and then solved pRF models for each vertex on the time-series averaged across all participants to compute group-average pRF maps on the *fsaverage* surface. The second and third methods differ in the order of operations: the second computes median parameters from all individual participants’ time series, and the third computes a single set of parameters on the averaged time series. These three implementations are described in detail below.

#### 2.6.1 Implementing the pRF model: Individual participants and the parameter-median data

First, we solved the pRF model for individual participants on their *fsnative* surface. For each participant, the time-series data across multiple preprocessed scans with the same stimulus were averaged to create a participant-averaged time-series. The participant-averaged time-series was then transformed to BOLD percent signal change (i.e., percent change at each TR from the mean signal across all TRs). We fit the pRF model to this BOLD signal in *fsnative* space. An example of the pRF model fit to an individual participant’s BOLD signal from a V1 and a V2 vertex is shown in **Figure 2A** and **B**.

**Figure 2.**
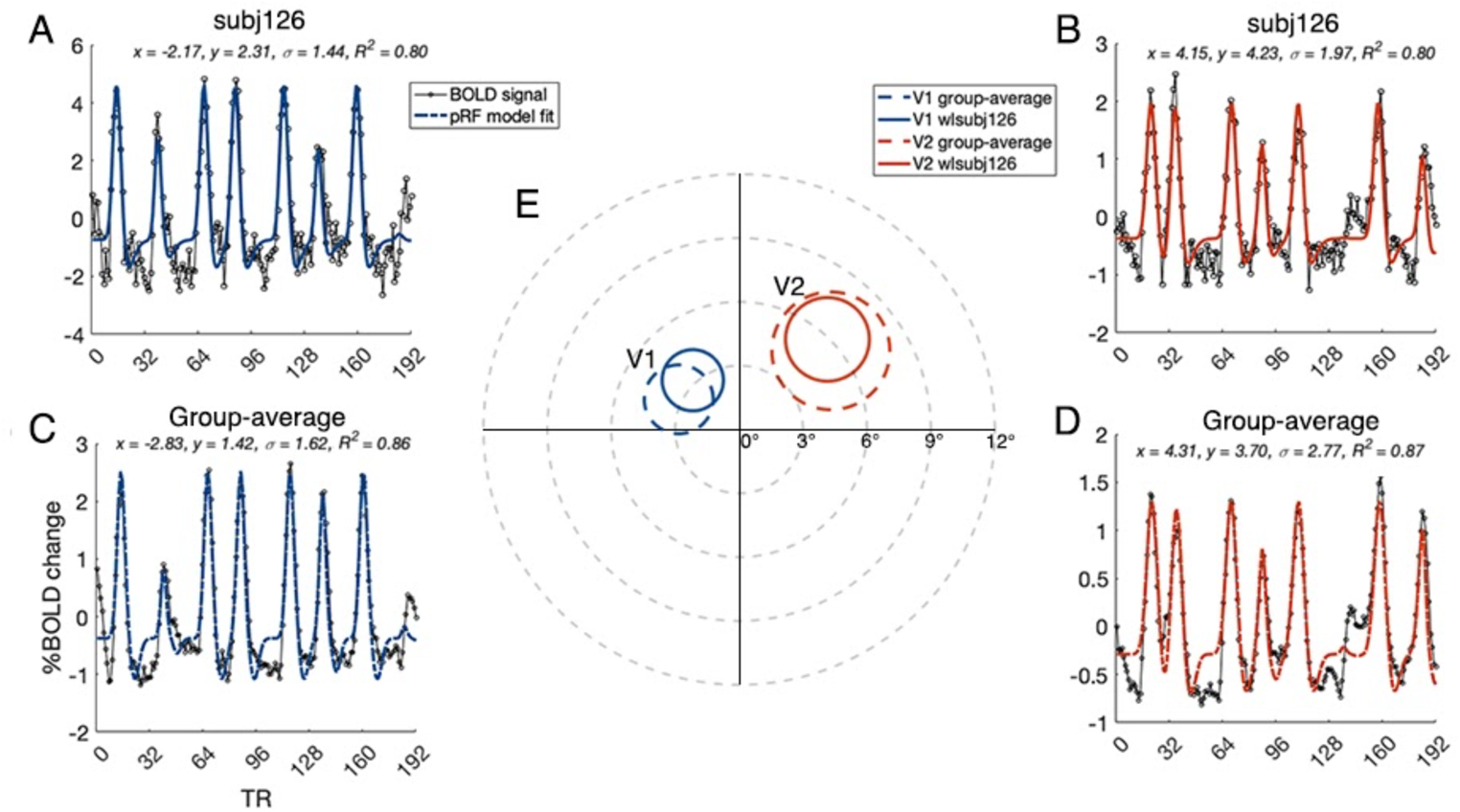
Examples of the pRF model fit to an individual participant’s (wlsubj126) and the group-average BOLD percent signal change for two example fsaverage vertices in V1 and V2. (**A-B**) The BOLD signal and pRF model fit for a V1 and V2 vertex for an individual participant. (**C-D**) The BOLD signal and pRF model fit for a V1 and V2 vertex for the group-average time-series data. (**E**) Visual field representation with pRFs plotted for V1 and V2. The inset text lists the predicted *x* and *y* values, representing the pRF center coordinates (in degrees), and the σ value, representing the pRF size (in degrees), and the R^2^ which is the variance explained of the pRF model fit to the BOLD signal.

Next, we computed parameter-median pRF maps. We computed the median (rather than the mean) as pRF estimates vary substantially across individuals (Harvey & Dumoulin, 2011; Song et al., 2015) and are not normally distributed (size and eccentricity are truncated at 0, and polar angle may be truncated near the vertical meridians). These parameter-median maps were used to complete vertex-wise comparisons between the NYU and HCP data. We used the FreeSurfer function *mri_surf2surf* to interpolate the pRF parameters by nearest neighbor from each participant’s *fsnative* surface onto the *fsaverage* surface. To generate the parameter-median polar angle and eccentricity maps, we calculated the median *x*, *y,* and *σ* values for each *fsaverage* vertex across all participants and converted these values to polar angle, eccentricity, and pRF size parameters.

#### 2.6.2 Implementing the pRF model: The group-average time-series data

We implemented the pRF model on group-averaged time-series data for map visualization and for visual comparison with the HCP data, which also include group-average time series and pRF model solutions (Benson et al, 2018).

We generated the group-average time-series as the vertex-wise average of each participants’ average time-series on the *fsaverage s*urface. To do so, we averaged each participants’ time-series across runs (number of runs varied between 4 and 12) on the *fsnative* surface. The participant average time-series was then resampled onto the *fsaverage* surface using nearest neighbor interpolation. Each of the participant-average time-series (n=44) were then averaged together to create a group-average time-series. The group-average time-series at each vertex was transformed into a group-average BOLD signal. The pRF model was then fit to the group-average BOLD signal to compute group-average pRF maps on the *fsaverage* surface. An example of the pRF model fit to the group-average BOLD signal from a V1 and a V2 vertex is shown in **Figure 2C** and **D**.

### 2.7 HCP 7T Retinotopy Dataset

The HCP 7T Retinotopy Dataset, as described by Benson et al. (2018), contains pRF model solutions for 181 participants. The methods for acquisition and analysis of these data are described in detail in the prior paper. We summarize a few key features here, particularly those that differ from the NYU data. The HCP data were collected at the Center for Magnetic Resonance Research at the University of Minnesota, using a Siemens 7T Magnetom scanner (1.6mm isotropic voxel size) and a 32-channel head coil. The data were processed using specific HCP pipelines, including multimodal surface matching (*’MSMall’*) for surface registration, as described by (Glasser et al., 2016). Notably, the publicly available HCP pRF estimates (https://osf.io/bw9ec/) were computed using the *analyzePRF* algorithm (Kay et al., 2013). Here, we recomputed the pRF estimates using *vistasoft* to remove any differences that might arise from differences in pRF software (see (Lage-Castellanos et al., 2020; Lerma-Usabiaga et al., 2020). We make these *vistasoft* pRF estimates publicly available via our OSF (Open Science Framework) repository (*link to be made available with publication*). These and other differences are summarized in **Table 1**.

**Table 1.** Key differences between the NYU and HCP Retinotopy Datasets.

|  | NYU | HCP |
| --- | --- | --- |
| **Participants** | 44 | 181 |
| **MRI Strength** | 3 Tesla | 7 Tesla |
| **Voxel Size** | 2mm isotropic | 1.6mm isotropic |
| **Stimulus Aperture** | Bars | Bars, wedges, and rings |
| **Stimulus Extent** | 12.4° (radius) | 8° (radius) |
| **Stimulus Width** | 3.1° | 2° |
| **Stimulus Update** | 3 Hz | 15 Hz |
| **Stimulus Motion** | Discrete steps | Continuous drift |
| **Pre-processing** | fMRIPrep | HCP Pipeline |
| **pRF Model** | *vistasoft* | *analyzePRF* for prior public release<br>(reanalyzed with *vistasoft* here) |

We implemented the *vistasoft* pRF model on two formats of HCP data. First, we implemented the pRF model on individual participants on the *fsaverage* surface. Using these data, we then computed HCP parameter-median pRF maps. Second, we implemented the pRF model on the group-average time-series on the *fsaverage* surface.

To implement the pRF model on individual HCP participants, we included six scans (2x bar runs, 2x wedge runs, and 2x expanding/contracting ring runs, at 5 mins each; see Benson et al. (2018) for examples of these stimuli). The time-series from these scans were then transformed to BOLD percent signal change and then concatenated together. The *vistasoft* pRF model was fit to this BOLD signal. We then computed HCP parmeter-median pRF maps to complete vertex-wise pRF comparisons with the NYU data; we calculated the median *x*, *y*, and *σ* values across all HCP participants and converted these values to polar angle, eccentricity, and pRF size parameters, for each *fsaverage* vertex.

To estimate the HCP group-average pRF maps, all 181 of the participants’ time-series on the *fsaverage* surface were averaged together at each vertex independently for each stimulus type (bar, wedge, and ring) to create group-average time-series. These were transformed to BOLD percent signal change at each vertex. We implemented the *vistasoft* pRF model on these HCP group-averaged BOLD signals to produce group-average pRF maps.

### 2.8 Defining visual area regions of interest

Regions of interest (ROIs) were defined using the Wang Maximum Probability Atlas (Wang et al., 2015). We identified V1, V2, V3, and hV4. The atlas defined ROIs were used for vertex-wise comparisons between the NYU and HCP Datasets. Additionally, we defined these ROIs by hand on the *fsnative* flatmaps for individual NYU participants using *Neuropythy* v0.11.9 (https://github.com/noahbenson/neuropythy; (Benson & Winawer, 2018), following the nomenclature of Wandell, Dumoulin, and Brewer (2007), and of Winawer & Witthoft (2015, 2017) for hV4. The ROI label files, and the code used to draw the ROIs, are made publicly available via the NYU Retinotopy Dataset OSF (*link to be made available with publication*). The hand-labeled V1 ROI was used to compute cortical magnification within the retinotopic maps.

### 2.9 Defining wedge-ROIs to measure polar-angle meridian asymmetries in cortical magnification

Recently, polar angle meridian asymmetries in V1 cortical magnification have been identified as retinotopic features in the HCP data (Benson et al., 2021). We tested whether these cortical magnification asymmetries are also found in the NYU Retinotopy Dataset. We defined a number of ROIs in V1 corresponding to wedges in the visual field. The wedges were defined using a combination of *Neuropythy* and custom MATLAB code.

The wedges varied in their polar angle center. They were centered on either the left or right horizontal meridian, the upper vertical meridian, or the lower vertical meridian. The polar angle extent of the wedge-width also varied. We tested a range of wedge widths: ±15°, ±25°, ±35°, ±45°, and ±55°. The surface area of the vertices within these wedge-ROIs was summed to find the amount of cortical surface dedicated to processing visual space within these wedges.

PRF polar angle estimates contain measurement noise. Thus, a wedge in the visual field does not typically project to a single enclosed region on the cortex, but rather to a patchy and dispersed set of regions (see Figure 8B in Benson & Winawer, 2018). This complicates the computation of surface area. It is likely that absent measurement noise, the ground truth retinotopic projection of a wedge in the visual field is a single, enclosed region on the cortical surface. Our goal in defining wedge-ROIs was to make reasonable estimates of the surface area of these cortical regions. To do so, we used a procedure similar to that described by (Benson et al., 2021). In brief, we define a cortical ROI corresponding to a wedge in visual space. We do so by generating several sub-ROIs, each of which is constrained to a narrow eccentricity band, and each of which has a width in the polar angle direction derived from the average location of a pool of vertices whose pRF coordinates lie near the wedge boundary in visual space. The eccentricity defined sub-ROIs are then concatenated to yield one full wedge. We describe the specific implementation in more detail below and provide a visualization of the process to define the iso-angle boundary of a 15° wedge extending from the upper vertical meridian in **Figure 3**.

**Figure 3.**
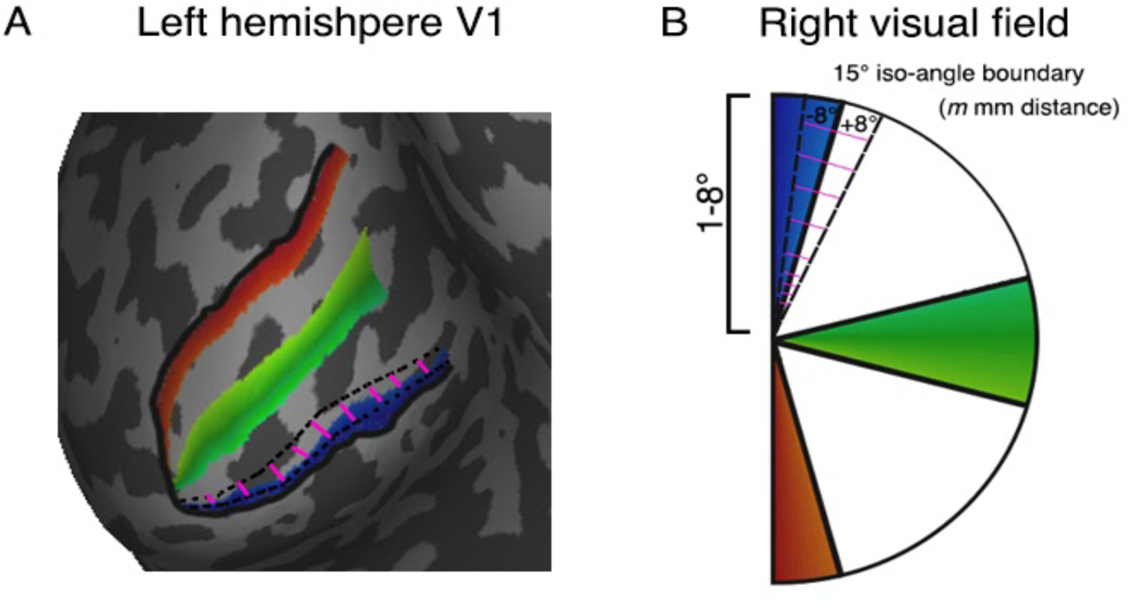
Defining wedge-ROIs and eccentricity band sub-ROIs. The wedge-ROI on the cortex (A) is defined as a wedge in the visual field (B). As there is measurement noise in the retinotopy data, the visual field wedge will not project to a single, enclosed region on the cortex. To circumvent this, we find all vertices in the V1 map which lie between 0 and *m* mm distance from the appropriate meridian. The vertices along the hand-drawn meridian are by definition at 0 mm from the meridian. To identify the distance *m_i_* of the +15° iso-angle boundary line from the upper vertical meridian in the i^th^ eccentricity bin, we calculate the average cortical distance of the vertices that fall ±8° around the 15° polar angle data. The ±8° boundaries are represented by the dashed black lines in (A) and (B). The average cortical distance of the +15° iso-angle boundary from the UVM is calculated separately for each of 10 log spaced bins (pink lines in A and B). We then define the wedge-ROI mask on the cortex (the blue data in A) as all the vertices between 0 mm distance and *m_i_* mm for each eccentricity sub-region. The cortical surface area within the wedge-ROI mask is summed across the 10 sub-regions. This is repeated for the right hemisphere (left visual field) to define the opposite portion of the UVM wedge.

**Figure 8.**
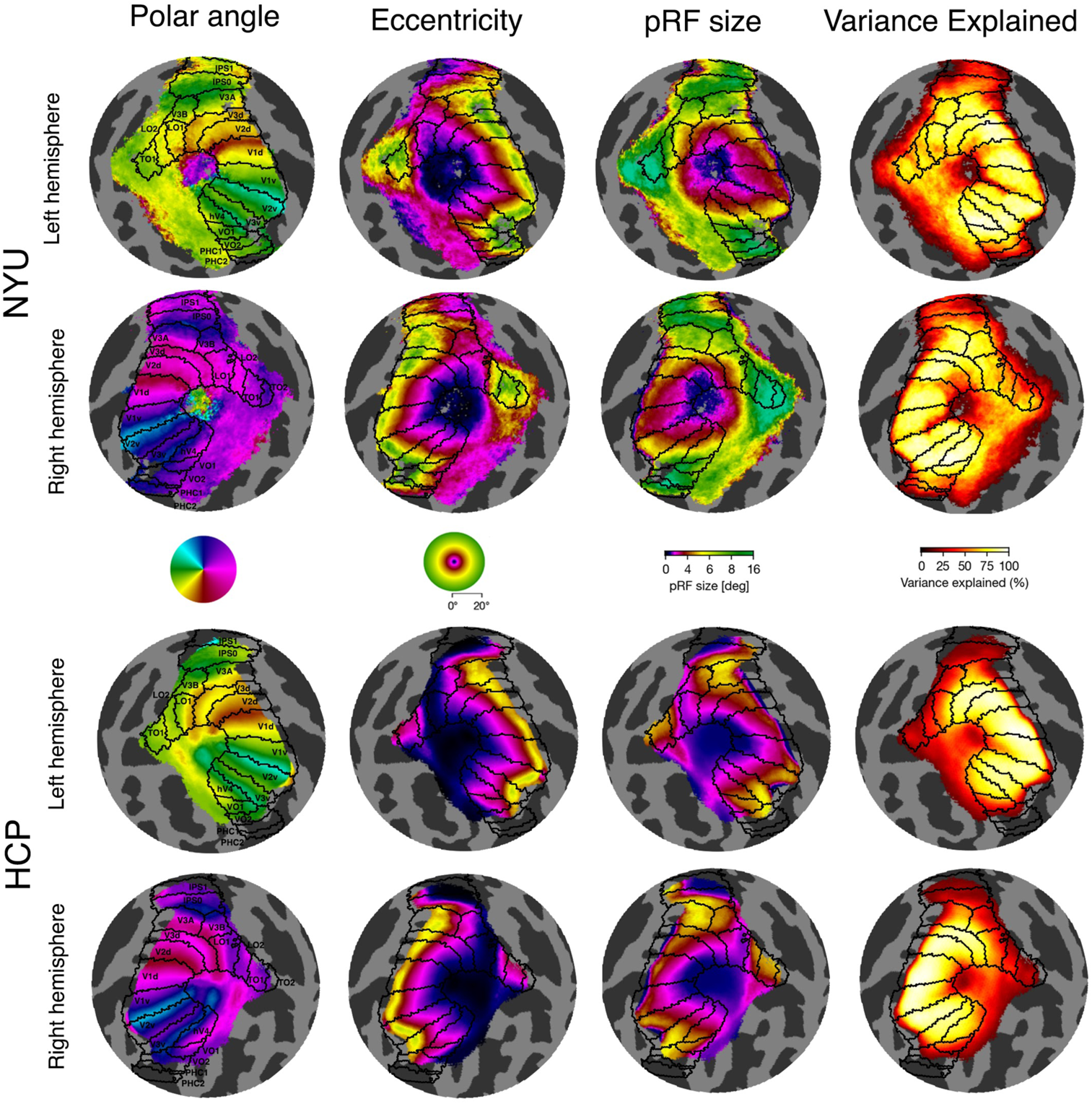
Median-parameter retinotopic maps for the NYU and HCP data shown on *fsaverage* flatmaps for the left and right hemisphere. The top two rows present the median-parameter maps for the NYU data, and the bottom two rows present the median-parameter maps for the HCP data. Black boundaries are ROI definitions derived from the Wang Maximum Probability Atlas (Wang, Mruczek, Arcaro, & Kastner, 2015). Note that pRF sizes are larger in the group-average time-series maps in Figure 7 than the parameter-average maps shown here; this is due to blurring of the fMRI signal during time-series averaging, resulting in larger pRF size estimates. Likewise, eccentricity estimates appear greater in the Figure 7 group-average time series maps; this is because of individual variation in the anterior V1 border across participants in the parameter-median data.

#### 2.9.1 Cortical distances

First, we computed the shortest distance on the cortical surface between each pair of vertices (using Dijkstra’s algorithm - the same algorithm that powers point-to-point navigation on google maps; (Dijkstra, 1959); and the appropriate cardinal meridian. For each participant and each hemisphere, we manually defined three line-ROIs in V1, for the horizontal, upper, and lower vertical meridians. The cortical representation of the horizontal meridian runs through the approximate middle of V1, the upper vertical meridian representation follows the V1/2 ventral boundary, and the lower vertical meridian representation follows the V1/2 dorsal boundary. The line-ROIs were drawn on the flattened cortical surface using *Neuropythy.* The manual definitions were informed by anatomy (curvature map) and pRF maps (polar angle and eccentricity). These line-ROIs were used to generate three *cortical distance maps* for each hemisphere. These cortical distance maps specify the distance of each vertex from each meridian.

#### 2.9.2 Eccentricity boundaries for sub-ROIs

Next, we divided the V1 map into 10 eccentricity bands, log spaced between 1° and 8° of eccentricity. These eccentricity bands were used as sub-ROIs. To ensure that each eccentricity band was a contiguous region, the bands were defined using the retinotopic maps generated by Bayesian inference (Benson & Winawer, 2018), which combines the participant’s vertex-wise pRF estimates with a previously defined retinotopic template to produce a denoised estimate of the visual field. The eccentricity bands were log-spaced in the visual field so that they would be approximately equally spaced in the cortex, as in Benson et al. (2021). The question we addressed here concerns asymmetries in surface area with respect to polar angle, thus, we did not rely on the Bayesian estimates of polar angle. The Bayesian optimization does not explicitly model cortical magnification and thus can induce noise in calculations of an ROI’s surface area. Instead, we used *Neuropythy* to implement an optimization that ‘cleaned’ polar angle pRF fits in V1 for each participant. This minimization seeks to adjust the pRF centers of the vertices as little as possible in order to simultaneously enforce a smooth cortical magnification map, as measured at the individual surface vertices, and correct the field-sign values across V1. Note that these Bayesian eccentricity estimates were used only for this ROI method, not for any comparisons of the pRF properties between datasets.

#### 2.9.3 Polar angle boundaries for sub-ROIs

Next, within each eccentricity band, we used the cortical distance maps to compute the average distance of an iso-angle line, representing the outer boundary of the wedge in the visual field, from the center of the wedge (and thus one of the visual field meridians). The iso-angle line was identified by calculating the average distance of the vertices, that had an R^2^ above 15%, in the region of cortex ‘around’ the iso-angle boundary at the defining polar angle value for the wedge-width (i.e., for a 15° wide wedge, this outer iso-angle boundary occurs at 15° in visual space). The vertices ‘around’ the iso-angle boundary were identified using the cleaned polar angle maps, and were defined as the vertices that fell ±8° polar angle from the iso-angle boundary. We identified the average cortical distance of these vertices to calculate the average cortical distance for the iso-angle line. This was repeated for each eccentricity band.

#### 2.9.4 Closing the sub-ROI

For each eccentricity band, we identified the vertices with a cortical distance between 0mm (i.e., those that fall along a meridian) and the average cortical distance of the iso-angle line. This process is repeated to create a mask for each eccentricity band, for each side of the wedge, and each meridian.

#### 2.9.5 Wedge-ROI

In the final step, we overlay the wedge mask on *cortical surface area maps.* These cortical surface area maps were generated for each participant using *Neuropythy* and specify the cortical surface area (in mm^2^) of each vertex on the *fsnative* surface. The cortical surface area is calculated by summing the surface area of the vertices within the wedge mask. The output value is the total surface area of the wedge. For each wedge, the surface areas from the left and right hemisphere are summed together to calculate the surface area for the full horizontal meridian, upper vertical meridian, and lower vertical meridian. The upper and lower vertical meridian surface areas are summed to find the surface area of the full vertical meridian.

### 2.10 Data visualization

To visualize individual participant retinotopic maps, we mapped each participant’s pRF data onto their inflated *fsnative* surface. The *fsnative* surface was then transformed into a sphere, and orthographically projected to a flatmap with pRF data rotated so that the occipital pole aligned to the center of the map (see **Figure 4** for these transformations). To visualize group-average time-series and parameter-average retinotopic maps, this process was repeated however the pRF data was mapped onto the *fsaverage* surface, rather than an *fsnative* surface.

**Figure 4.**
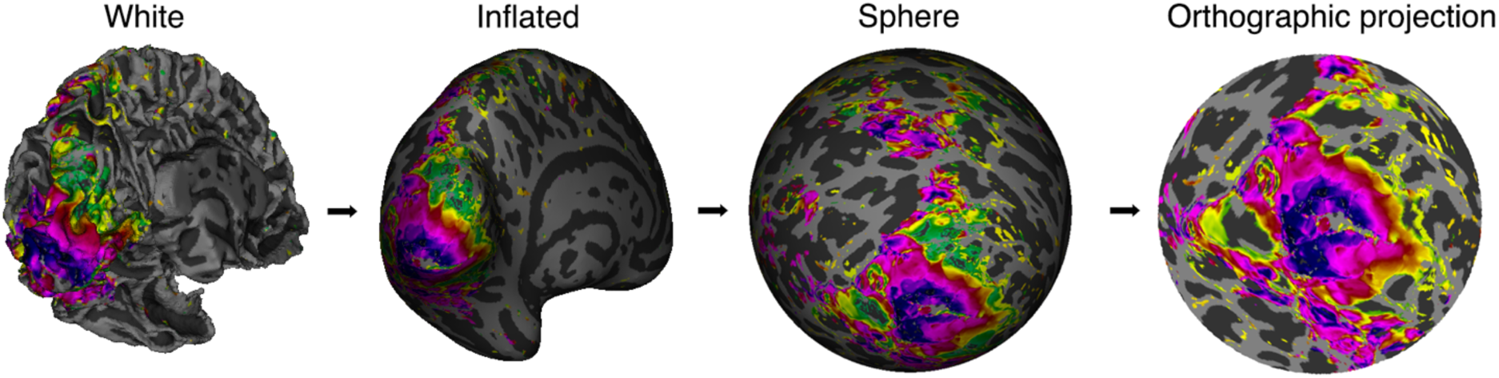
Transform of a left hemisphere cortical surface from white matter mesh through to orthographic projection on a flatmap. The pRF data (in this example, subj004’s eccentricity map) are projected onto the *fsnative* surface. The surface is then inflated and smoothed, warped into a sphere, and then rendered as a flatmap with the occipital pole rotated to be at the center of the map.

## 3. Results

We present our results across three sections. First, because the NYU Retinotopy Dataset is new and publicly available (access via: *link to be made available with publication*), we provide a description, including visualizations of pRF maps computed from individual participants and the group-average time-series. This communicates the quality and consistency of the data, similarities and differences among individual maps and group-average maps, the effect of how the average is computed, and the agreement of our data to a previously published atlas. In addition, we provide publicly available Jupyter Notebook code to visualize individual retinotopic maps, via cortical flatmap or inflated mesh, for all of the NYU and HCP data. Second, we quantify the similarity of NYU and HCP vertex-wise pRF estimates (polar angle, eccentricity, and pRF size) in V1, V2, V3, and hV4. Third, we report whether the polar angle asymmetries in V1 cortical magnification previously reported in the HCP data (Benson et al., 2021) can be generalized to the NYU data.

### 3.1 NYU Retinotopy Dataset: Individual participants

PRF model solutions were computed for each of the 44 individual participants on the *fsnative* surface. Examples of polar angle, eccentricity, and pRF size maps for three example participants (subj004, subj014, and subj042) are illustrated in **Figure 5**. The data are plotted similar to Figure 7 in Benson et al. (2018) for ease of comparison to the HCP retinotopy single-participant data. V1, V2, V3, and hV4 ROIs are drawn in black along the polar angle boundaries. The same pRF maps on inflated meshes are available in **Supplementary Materials Figure S1**. The retinotopic maps are visualized out to 20° eccentricity to show the full data available; it is possible to find pRFs centered outside the maximum stimulus extent (12.4°), as some portion of the neural receptive fields will overlap the pRF stimulus, thereby allowing one to estimate their pRF parameters. We do not include the pRFs beyond the stimulus extent in our analyses as they may be biased towards lower eccentricities as the same response might be explained by either a small pRF close to the stimulus edge or a large pRF far from it.

**Figure 5.**
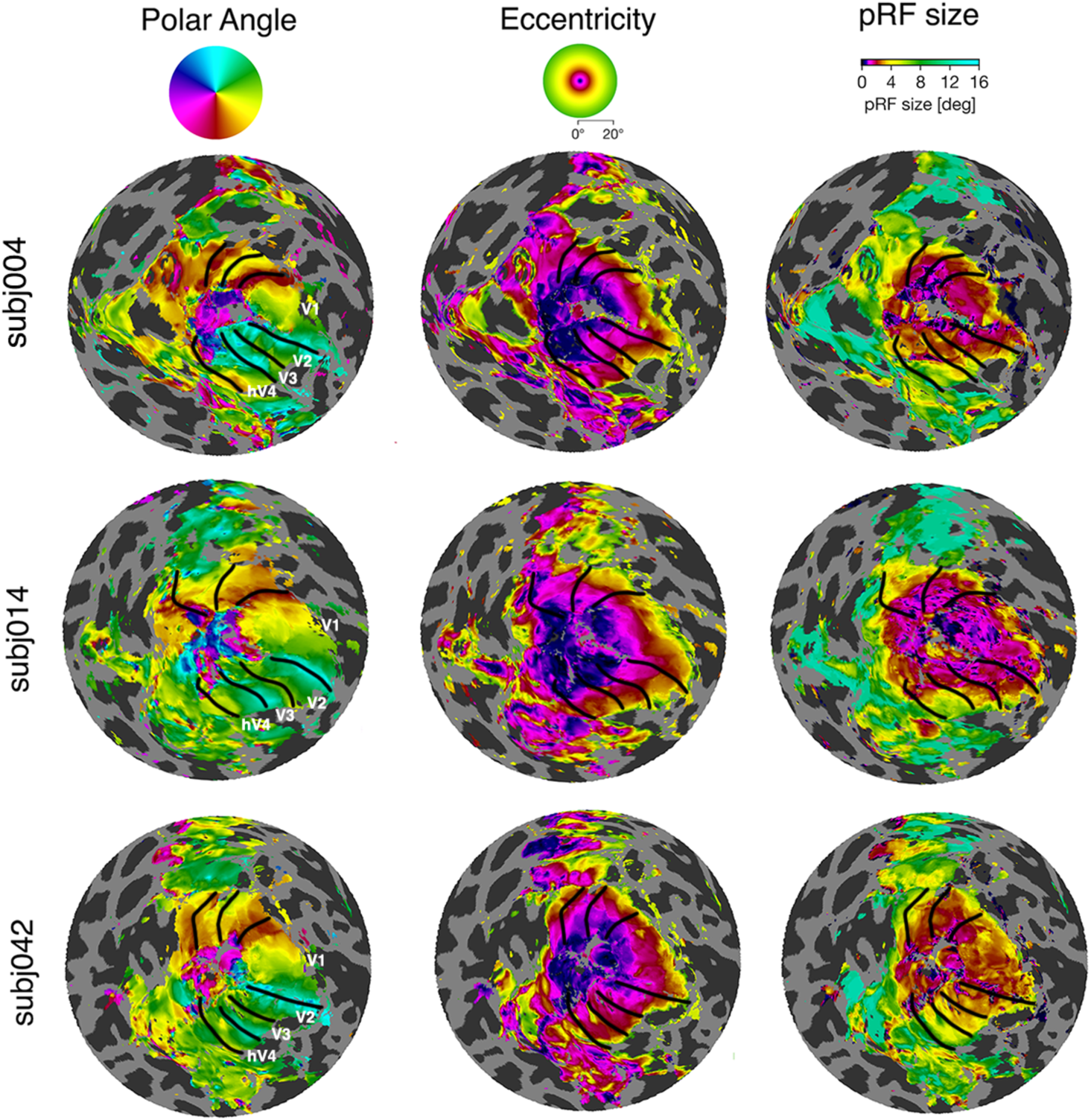
Examples of polar angle, eccentricity, and pRF size maps for three individual participants (subj004, subj014, and subj042). The maps are presented on cortical flat maps on the *fsnative* surface of the left hemisphere. V1, V2, V3, and hV4 ROI boundaries are defined in black. R^2^ threshold = > .12.

**Figure 7.**
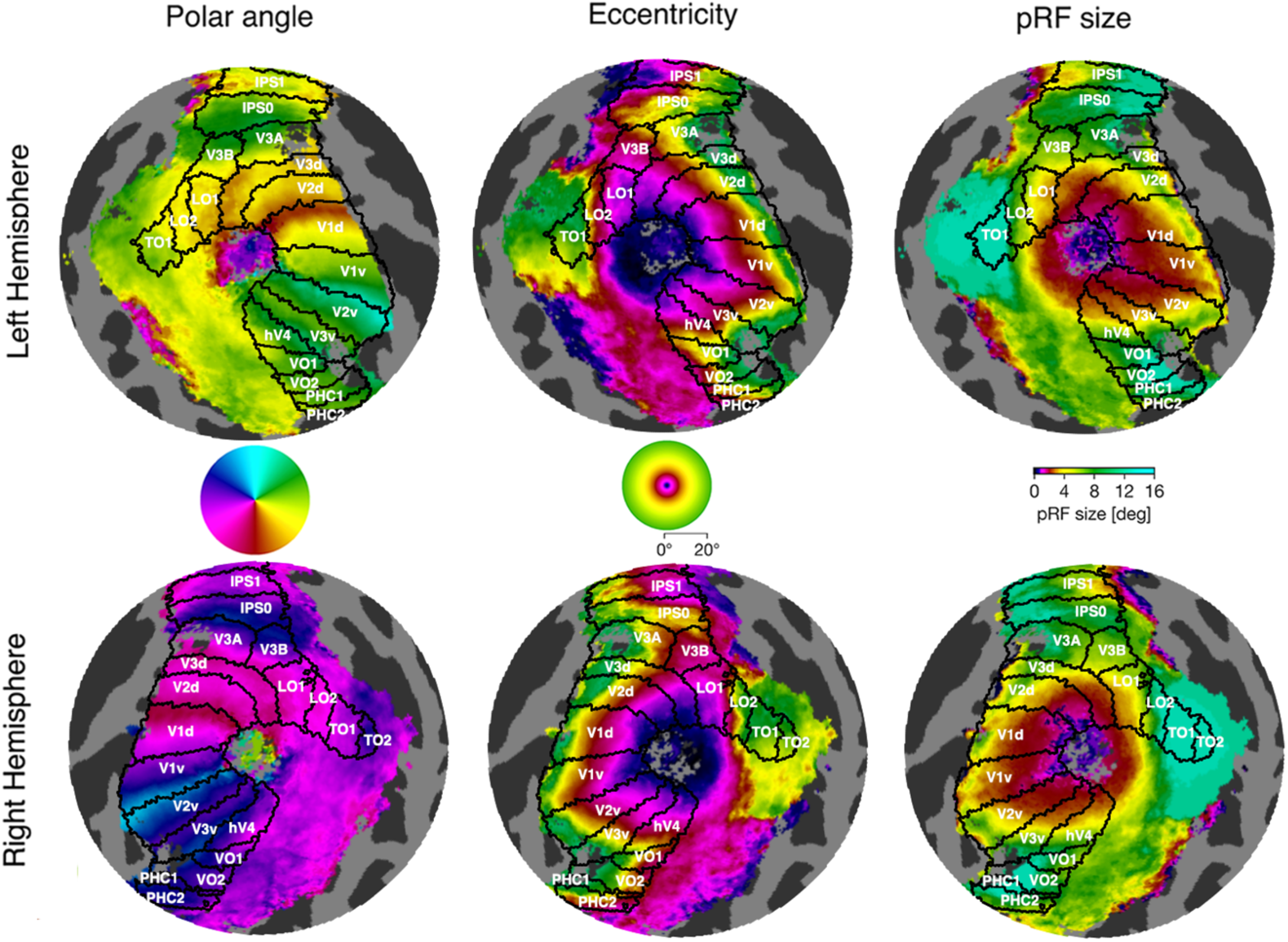
Group-average NYU retinotopic maps shown on the *fsaverage* flatmaps for the left and right hemisphere. The top row presents flatmaps of the left hemisphere (representing the right visual hemifield) and the bottom row presents flatmaps from the right hemisphere (representing the left visual hemifield). Black boundaries are ROI definitions derived from the Wang Maximum Probability Atlas (Wang, Mruczek, Arcaro, & Kastner, 2015).

The individual participant maps show clear polar angle, eccentricity, and pRF size representations, and for each participant the maps follow the same broad retinotopic organization, though they differ in detail. For example, the location and size of dorsal V3 are quite different between subj004 and subj014. In addition, the representation of the lower visual field in hV4 differs, with subj004 but not the other two subjects showing representation all the way to the lower vertical meridian (red) (**Figure 5**).

### 3.2 NYU Retinotopy Dataset: Group-average time-series data

We averaged across 44 participant-averaged time-series on the *fsaverage* surface to create a group-average time-series. The group-average time-series will blur out some features of the fMRI response, particularly in higher-level visual areas where the topography of the maps might not be closely linked to the sulcal patterning. In V1, V2, and V3, many features of the retinotopic maps are preserved after averaging, and the time-averaged data offer a succinct summary of the dataset with a high signal-to-noise ratio. The HCP Retinotopy Dataset also includes the group-averaged time-series data, and hence doing so makes for a useful comparison.

The group-average time-series was transformed to BOLD signal and pRF model solutions were calculated. In **Figure 6**, we present retinotopic maps of the group-averaged data projected onto the inflated *fsaverage* surface for the left and right hemispheres. The visual field representations are located across the occipital cortex and extend along dorsal and ventral regions, as well as frontoparietal cortex (Mackey et al., 2017). These visualizations highlight large-scale structure across cortex, such as the continuous parafoveal representation (magenta) from the parietal lobe to the occipital lobe to the ventral surface. However, the inflated surfaces self-occlude, so we next show flattened views of the two hemispheres separately.

**Figure 6.**
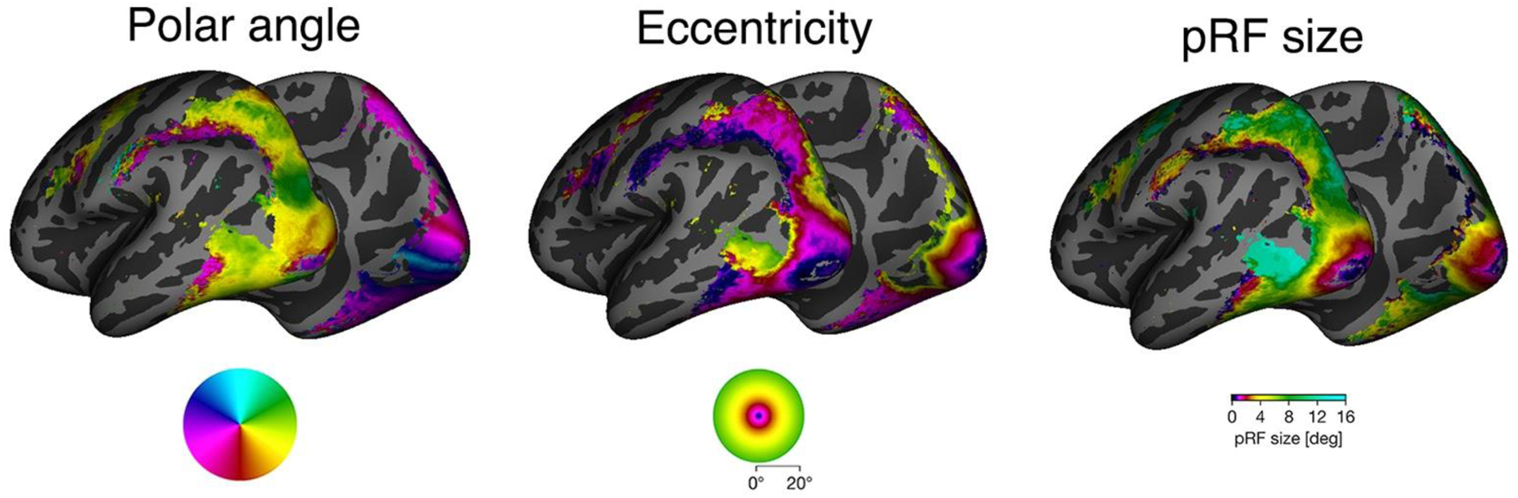
Group-average time-series polar angle, eccentricity, and pRF size maps shown on the left and right hemisphere *fsaverage* inflated surface.

In **Figure 7** we present the group-average polar angle, eccentricity, and pRF size maps for the left and right hemispheres on an *fsaverage* flatmaps, with the occipital pole aligned at the center of the flatmap. The Wang Maximum Probability Atlas (Wang et al., 2015) is overlaid on the retinotopic maps. The ROI borders from the Wang Atlas are well-aligned to the polar angle reversals. However, the V1/V2 dorsal boundary appears to be slightly misaligned, falling slightly superior to the red polar angle data. This is common in both the left and right hemisphere.

Thus, we suspect that the vertices falling within the Wang V1d ROI contain a small number of vertices from the V2d ROI.

The maps show the expected retinotopic organization, particularly in the early visual areas; there is a clear polar angle representation with reversals occurring at the ROI boundaries, eccentricity estimates increase with distance from the foveal representation, and pRF sizes increase with eccentricity while scaling to be larger in higher-order areas.

There is an artifact in polar angle estimates around the occipital pole, representing the most foveal region of the visual field. These regions show anomalous polar angle representations of the ipsilateral visual field. For example, in **Figure 7**, there is a patch of purple and blue polar angle data around the fovea of the left hemisphere, and there is a patch of yellow and green polar angle data for the right hemisphere. A similar effect was found for individual participants (see **Figure 5**); however, it was not found in the HCP group-average time-series data, perhaps due to the inclusion of rotating wedge stimuli in the HCP experiments. (See **Supplementary Materials S2** for HCP group-average time-series pRF maps from *vistasoft*).

### 3.3 Vertex-wise reproducibility of pRF parameters between the NYU and HCP retinotopy data: eccentricity, polar angle, and pRF size

We tested the similarity of polar angle, eccentricity, and pRF size estimates between the NYU and the HCP Retinotopy Datasets. To compare the two datasets, we identified the median pRF parameters at each vertex on the *fsaverage* surface separately for the two datasets. This puts both datasets on a standardized and common surface, making it possible to complete vertex-wise comparisons. For comparison, we group the vertices using V1, V2, V3, and hV4 ROIs from the Wang Atlas. The Wang Atlas contains ROIs that are originally defined on neither the HCP or the NYU data. This provides an automated, reproducible way to bin data in the same manner for the two datasets, even though the atlas-based ROIs only approximately match the visual field maps defined at the level of individual participants. Visualizations of the NYU and HCP parameter-median pRF maps, including variance explained maps, are presented **Figure 8**.

We compared pRF estimates for vertices that passed inclusion criteria within each ROI (V1, V2, V3, and hV4). Data close to the fovea are challenging to measure, especially for the NYU data which show ipsilateral polar angle representations around the fovea. Conversely, potential edge effects occur at higher eccentricities in the HCP data, where pRF size begins to decrease with increasing eccentricity beyond 6°. Thus, we retained vertices that fell between 0.2° and 6° of eccentricity and survived a 40% variance explained threshold. A vertex was required to meet these criteria for both the NYU and HCP parameter-median data to be included in the analysis. The variance explained threshold removed 14.4% of the vertices within this eccentricity range in the V1 to hV4 maps. In **Figure 9**, we present scatterplots comparing vertex-wise NYU and HCP pRF estimates for vertices within V1, V2, V3, and hV4.

**Figure 9.**
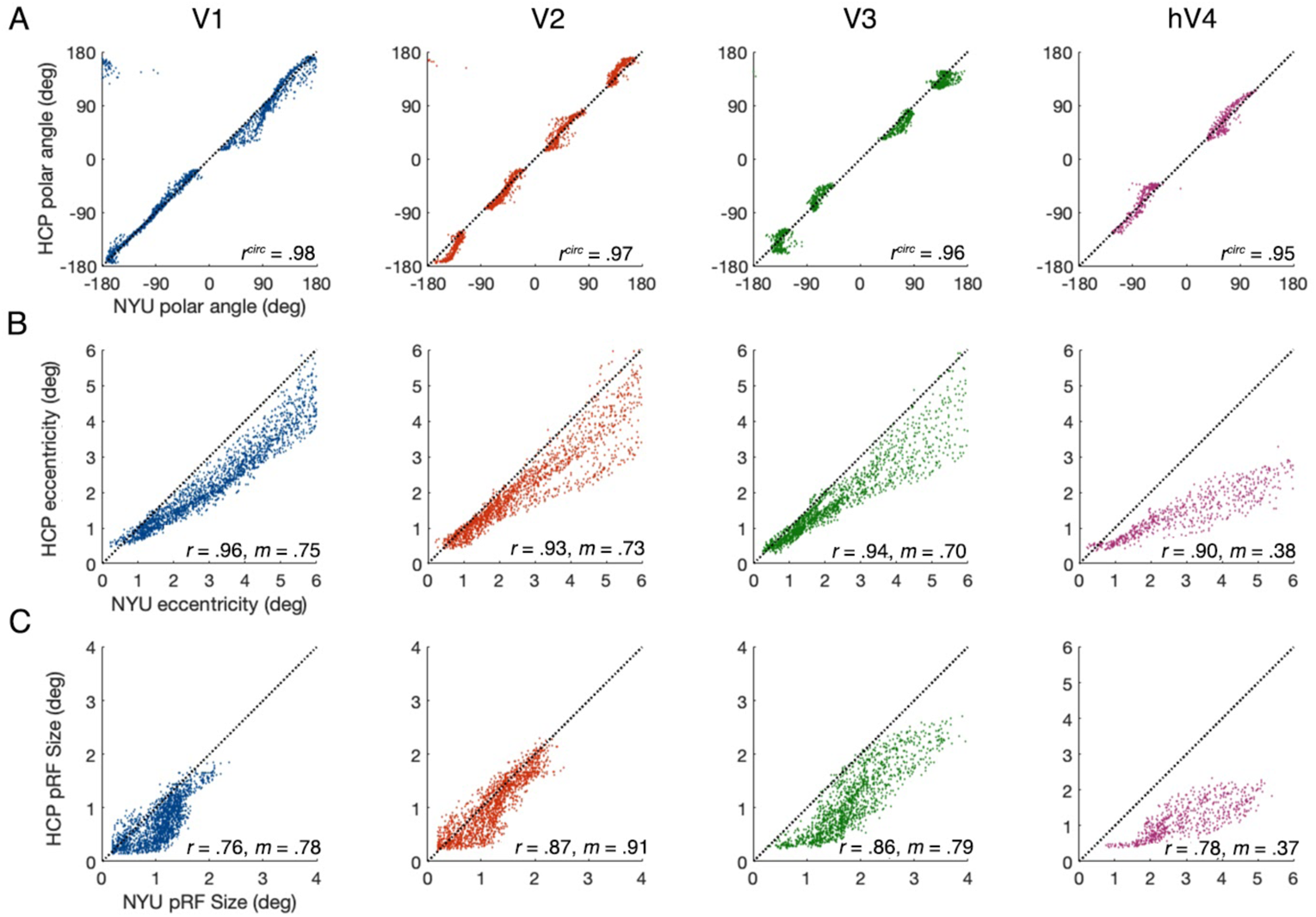
Scatterplots comparing vertex-wise pRF estimates from the NYU and HCP parameter-median data (visualized in Figure 8) within V1, V2, V3, and hV4. Each data point represents an *fsaverage* vertex that has survived a double mask. (**A**) Vertex-wise comparison of polar angle estimates. 0° to +180° polar angle represents the right visual hemifield (with 0° at the upper vertical meridian and −180° the lower vertical meridian), and 0° to −180° polar angle represents the left visual hemifield. (**B**) Vertex-wise comparison of NYU and HCP eccentricity estimates. **(C)** Vertex-wise comparison of NYU and HCP pRF size estimates. Note the different values of the X and Y axis for the hV4 plot in panel C. The black dashed line represents *y* = *x*. All reported *r* values are highly significant (*p* < .001) and *m* represents slope.

The polar angle estimates fall close to the identity line and are highly correlated in V1 (circular correlations (Fisher & Lee, 1983); *r^circ^* = .98), V2 (*r^circ^* = .97), V3 (*r^circ^* = .96), and hV4 (*r^circ^* = .95), indicating a close match between NYU and HCP polar angle data (**Figure 9A**). Notably, there are gaps in the data along the polar angle reversals for each visual field map (0° and ±180° for V1 and hV4, and 0°, ±90, and ±180° for V2 and V3). This is likely due to averaging effects, both at the level of individual participants, because voxels pool signals from many neurons, and at the group level, because the borders between visual areas are not in perfect register across participants.

The eccentricity estimates are also highly correlated in V1 (linear correlations; *r* = .96), V2 (*r* = .93), V3 (*r* = .94), and hV4 (*r* = .90) (**Figure 9B**). However, unlike polar angle estimates, there are systematic biases in eccentricity. The NYU eccentricity estimates are generally greater than those from the HCP data in all ROIs. The slopes of the data were similar in V1 (*m* = .75), V2 (*m* = .73), V3 (*m* = .70), and lower in hV4 (*m* = .38). Thus, in V1 to V3, for every 1° change in NYU eccentricity, the HCP eccentricity estimate changes by ~0.70°. The larger disagreement between the two datasets in hV4 is addressed in the Discussion. Comparisons of NYU and HCP cortical magnification plotted as a function of eccentricity (V1-V3) are available in **Figure S3**.

The pRF size estimates are well correlated in V1 (linear correlations; *r* = .76), V2 (*r* = .87), V3 (*r* = .86), and hV4 (*r* = .78) (**Figure 9C**). Similar to the eccentricity estimates, there are systematic biases in pRF size: the NYU pRF size estimates are typically larger than those derived from the HCP data. The slopes of the data are similar in V1 (*m* = .78), V2 (*m* = .91), V3 (*m* = .79), and as with eccentricity, lower in hV4 (*m* = .37).

Typically, fMRI studies use far fewer participants than either of these two datasets. An important question is how reproducible the data would be for smaller datasets. To answer this, we repeated the correlation analyses for small subsamples (20 participants from each dataset, with replacement), repeated 500 times (**Figure S4**). The correlations remain high, even with a reduction in sample size.

### 3.4 PRF size vs eccentricity

The previous analysis showed differences between the two datasets in pRF size and eccentricity for the vertex-wise comparisons. Rather than comparing parameters matched by cortical location, we compared parameters matched by eccentricity. Here, we plot the NYU and HCP pRF sizes as a function of eccentricity for V1, V2, V3, and hV4, and fit a linear function through the data. The NYU data (solid lines) and HCP data (dotted lines) show an increase in pRF size as a function of eccentricity and the pRF sizes scale to be larger in sequentially higher order visual areas for each dataset (**Figure 10**). When plotting the pRF sizes relative to eccentricity, the pRF sizes appear more similar between the two datasets than they do when comparing the datasets aligned by location (**Figure 9C**). This is particularly so for V1 to V3. Nonetheless, even when plotted as a function of eccentricity, some differences between the two datasets remain. Specifically, the HCP data converge closer to the origin than the NYU data, and the HCP hV4 data are generally smaller than NYU hV4 data. See Supplementary Materials **Figures S6 and S9** for analysis comparing parameter-median pRF estimates for the HCP all stimuli data against the HCP bar only data, and Supplementary Materials **Figures S7 and S8** for analysis comparing parameter-median pRF estimates for the NYU data against HCP bar only data.

**Figure 10.**
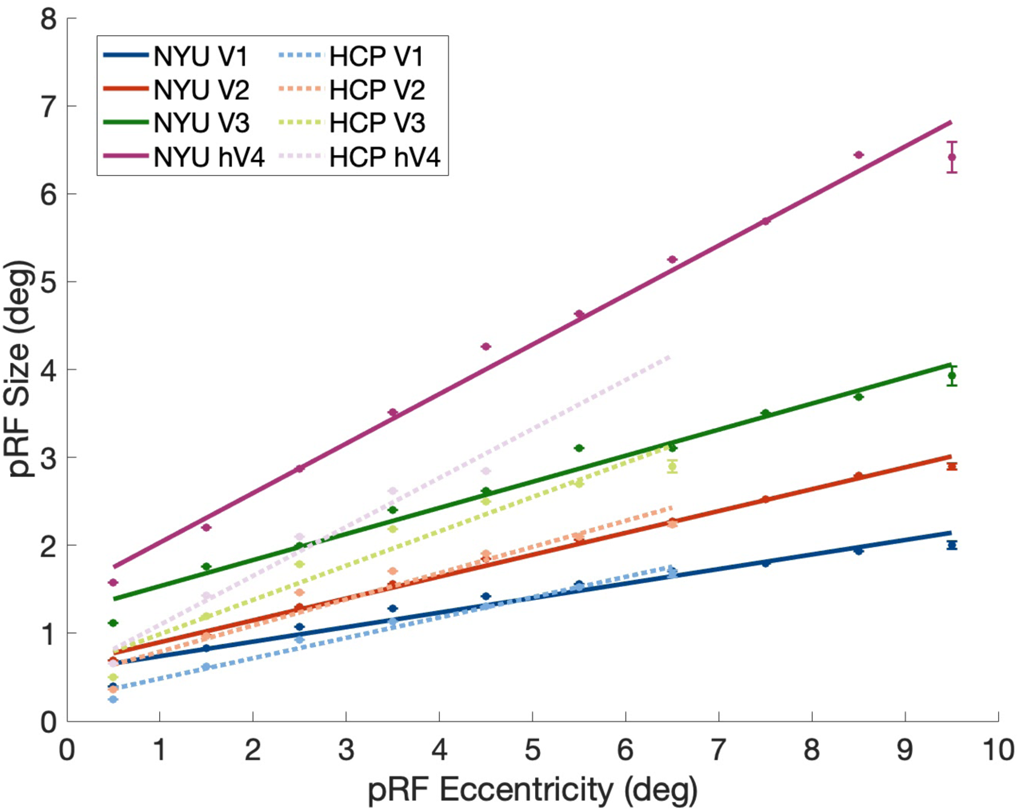
PRF sizes as a function of eccentricity for the NYU (solid lines) and HCP (dashed lines) parameter-median data for V1, V2, V3, and hV4. pRF sizes are binned into 1° eccentricity bins and a linear function is fit to the mean sizes. The NYU data are plotted to 0.5°-9.5° eccentricity, whereas the HCP data are plotted out to 0.5°-6.5° eccentricity due to differences in maximum stimulus extent. Note that the lines do not converge to the origin. If they are extended, then the x-axis intercepts for the NYU fits are: V1: −3.5°, V2: −2.6°, V3: −4.2°, and hV4: −2.6°. The x-axis intercepts for the HCP fits are: V1: −1.1°, V2: −1.7°, V3: −1.5°, and hV4: −1.0°. Error bars are ±1 SEM (vertices per bin), R^2^ threshold => .40.

### 3.5 How accurately do the NYU pRF parameters predict the HCP BOLD signal?

One advantage of the pRF model is that it is generative. Once the pRF parameters for a vertex are estimated, one can predict the complete time-series to other stimuli and experiments. The stimulus sequence of the NYU experiment differed from the stimulus sequence used in the HCP experiment in many ways. The HCP experiments included wedges and rings, which the NYU experiments did not; there were numerous differences just for the bar stimuli alone used in both experiments, including the maximal stimulus extent, discrete steps vs continuous motion, the speed of the stimulus sweeps across the visual field, whether diagonal sweeps were included, etc. A good and reliable computational model should be able to predict fMRI responses to other stimuli in other circumstances. One test of the quality of the NYU Retinotopy data, and of the pRF model itself, is how well the NYU pRF models can explain the BOLD time-series from a separate dataset that uses different stimulus types and has been collected under different circumstances, such as the HCP Retinotopy data. Further, there is a non-linear relation between predicted pRF parameters and the BOLD signal. A small change in a pRF parameter can cause a large change in the predicted time-series, and vice versa. Thus, one cannot directly infer how well the time-series (or predicted time-series) will match based on comparing the pRF parameters alone.

Thus, we tested how accurately the NYU pRF estimates, together with the HPC stimulus, can predict the HCP BOLD time-series. Specifically, we multiplied the NYU pRFs by the HCP stimulus aperture sequence at each time point (after convolving the stimulus with an HRF derived from the HCP data). For each vertex, given that the gain differs due to field strength and other factors, we then applied a multiplicative and additive term to best fit the HCP BOLD signal. From this we calculated the variance explained value, which provides a quantification of how well a predicted time-series derived from the NYU pRF parameters can explain the HCP BOLD signal. This measure has the merit of reducing a multiple parameter comparison (polar angle, eccentricity, and pRF size) to a single summary value per vertex.

In **Figure 11A** we present variance explained maps on the left and right hemispheres across the full surface of the brain. These maps show how well the predicted time-series generated from NYU pRF parameters can be fit to the HCP BOLD signal, thus the pRF parameters are fit to different data than they are evaluated on (except for the different scale factors of the time-series). The variance explained of the fits is high (above 70%) within V1-hV4 and remains high in extended ventral and dorsal regions of visual cortex.

**Figure 11.**
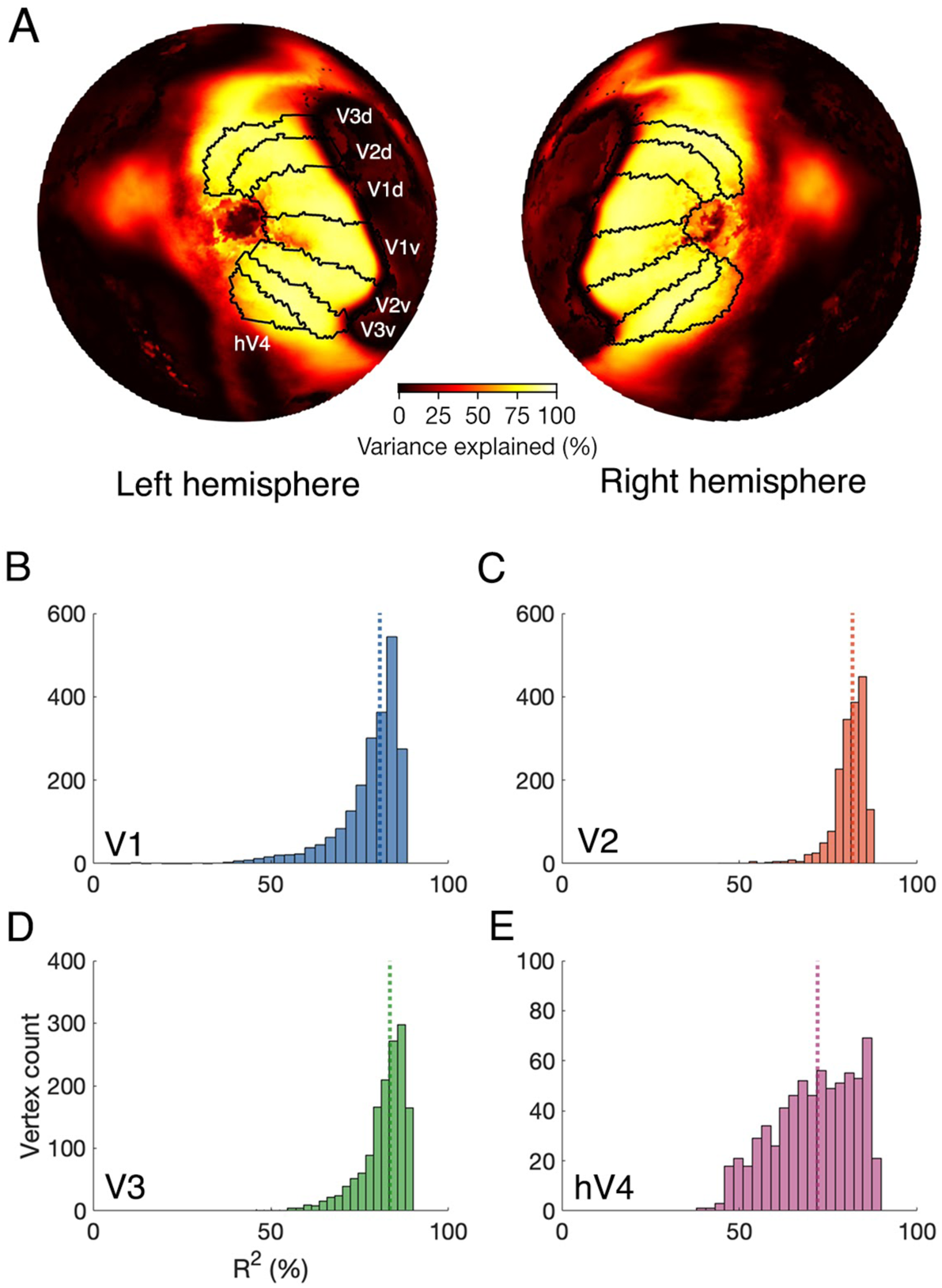
Variance explained maps of the NYU-derived predicted time-series fit to the HCP BOLD signal for the full left and right hemisphere, and histograms of the variance explained of the NYU-derived predicted time-series fits in V1, V2, V3, and hV4. **(A)** *Fsaverage* flatmaps illustrating the variance explained of the recreated NYU-derived model predictions fit to the HCP BOLD signal. **(B-E)** Histograms of the vertex-wise variance explained (%) of the NYU-derived model fits for V1 to hV4. The median variance explained of the fits are shown as a dotted colored vertical line. Data are restricted to V1 to hV4 defined by the Wang atlas and between 0.2 - 6° eccentricity.

The median variance explained of NYU-derived predicted time-series fit to the HCP BOLD signal was: 80% for V1 vertices, 82% for V2 vertices, 84% for V3 vertices, and 72% for hV4 vertices (**Figure 11B-E**). The NYU-derived predicted time-series can be used to explain the HCP BOLD signal, even though the parameters used to build the recreated NYU predicted time-series come from a different experiment that includes only bar stimuli (whereas the HCP BOLD signal is elicited from bars, wedges, and rings).

### 3.6 Polar-angle meridian asymmetries in V1 cortical magnification

Recent work has identified polar angle asymmetries in V1 cortical magnification using the HCP 7T Retinotopy Dataset. More cortical surface area is dedicated to processing the horizontal than vertical visual field meridian (a cortical HVA), and to the lower vertical than the upper vertical meridian (a cortical VMA) (Benson et al., 2021; Silva et al., 2018). The surface area as a function of polar angle aligns with performance on psychophysical tasks (Benson et al., 2021). Here, we tested whether these cortical surface area asymmetries generalize to the NYU Retinotopy Dataset.

For each participant, we calculated the cortical surface area of wedges that were centered on the horizontal, upper vertical, and lower vertical visual field meridians (see **Figure 12A** for an example). The wedges increased in width, from ±15° to ±55°, and spanned 1° to 8° of eccentricity.

**Figure 12.**
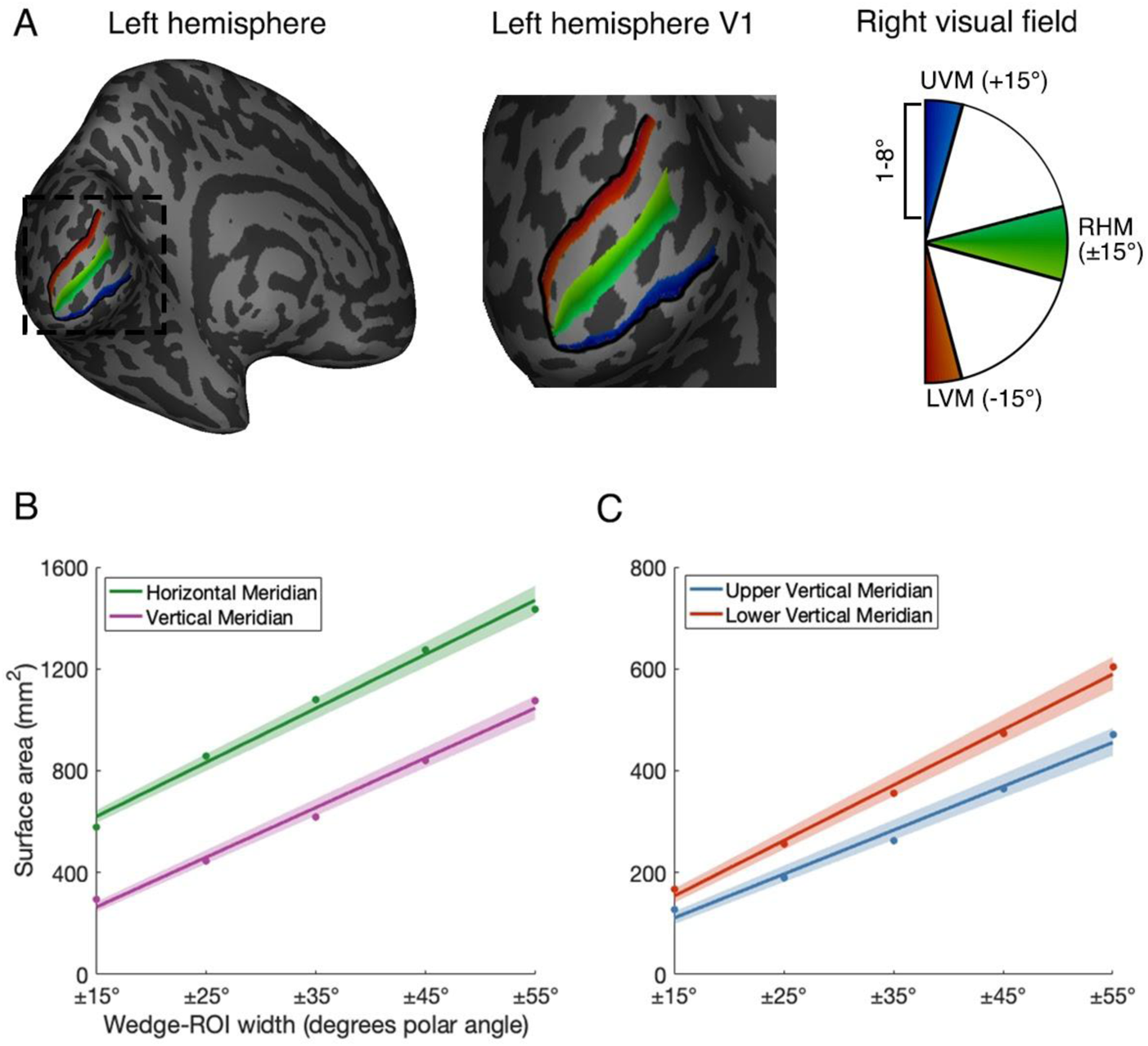
Examples of wedges in the visual field and their representation on the cortical surface of V1, and group-mean cortical surface area measurements plotted as a function of wedge width. **(A)** An inflated left hemisphere. The black borders show the boundaries of V1. The right visual field is mapped onto the left hemisphere of V1, and the wedges are centered on the upper vertical meridian, right horizontal meridian, and lower vertical meridian. We sum the cortical surface area of the vertices within each colored wedge to find the total surface area of each wedge. The horizontal meridian wedge is ±15° whereas the upper vertical meridian and lower vertical meridian are +15° and −15° because the full upper vertical meridian and lower vertical meridian wedge is formed by summing cortical space from the left *and* right hemisphere. **(B)** Cortical surface area measurements for wedges centered on the horizontal and vertical meridians, plotted as a function of wedge-width. **(C)** Cortical surface area measurements for wedges centered on the upper and lower vertical meridians, plotted as a function of wedge-width. Colored lines represent the average of 1000 bootstrapped linear fits to the data (colored data points). The shaded error bar represents the 68% bootstrapped CI of the linear fit to the data.

We were able to confirm the cortical HVA and VMA in our data. In **Figure 12B** and **12C**, we present the mean V1 surface area measurements for the wedges. In agreement with a recent study (Benson et al., 2021), there is more cortical surface dedicated to processing the horizontal than the vertical meridian (**Figure 12B**). Similarly, there is more cortical surface area dedicated to processing the lower vertical than the upper vertical meridian (**Figure 12C**).

Next, we calculated HVA and VMA asymmetry indices for each participant and each wedge. The HVA asymmetry index was calculated as the difference in surface area between the vertical meridian and the horizontal meridian, divided by the mean of the two, multiplied by 100. The VMA asymmetry index was calculated as the difference in surface area between the upper vertical meridian and the lower vertical meridian, divided by the mean of the two, multiplied by 100. Thus, a positive HVA or VMA asymmetry indicates an effect in line with the behavioral HVA or VMA and a larger index represents a stronger asymmetry.

The HVA asymmetry (**Figure 13A**) was around 65% for a ±15° wedge and the asymmetry decreased with increasing wedge-width. Thus, near the meridians, there is almost twice as much surface area dedicated to processing the horizontal compared to the vertical visual field meridian. The VMA (**Figure 13B**) was smaller, though still substantial, with around a 30% asymmetry for a ±15° wedge, and again, the asymmetry decreased with increasing wedge-width. For both the HVA and VMA, as the wedge width increases and the asymmetry decreases, the confidence interval increases. Hence, the most robust asymmetries for wedge-ROIs are close to the meridians. HVA and VMA asymmetries for incremental ±5° wedge are reported in **Supplementary Materials Figure S5**.

**Figure 13.**
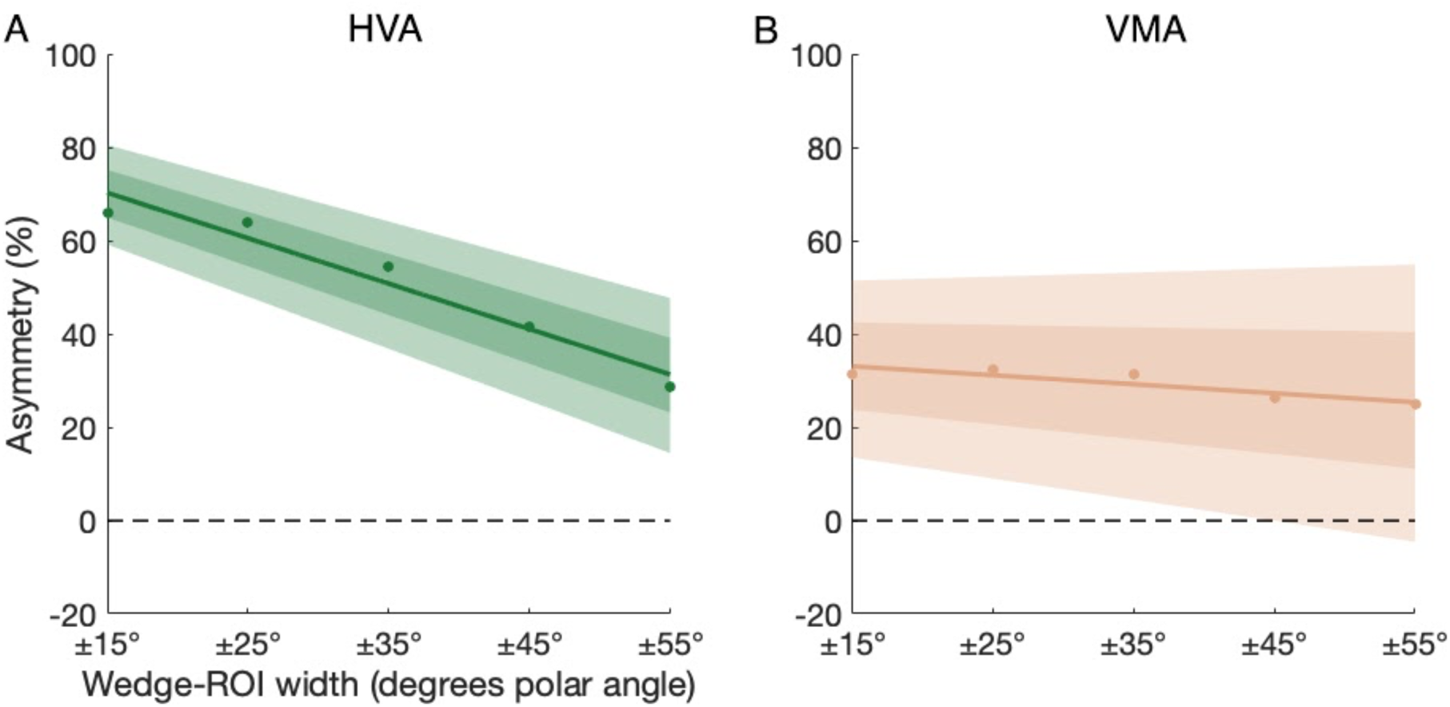
Group-mean HVA and VMA indices plotted as a function of wedge width. Green and orange lines represent the average of 1000 bootstrapped linear fits to the mean values (colored data points) at each wedge width. The light shaded error bar represents the 95% bootstrapped CI and the dark shaded error bar represents the 68% bootstrapped CI.

## 4. Discussion

Here, we have described a new, publicly available dataset; the NYU Retinotopy Dataset. We then tested the cross-dataset reproducibility between the NYU and HCP Retinotopy Datasets. Although these two datasets differ in many aspects, our goal was to examine how similar they are in terms of the retinotopic properties of human V1 through to hV4. First, we tested the reproducibility of vertex-wise pRF estimates in V1, V2, V3, and hV4. We found them to be strongly correlated between the two datasets, but with systematically larger eccentricities and pRF sizes in the NYU data. Second, we tested how well a predicted time-series derived from the NYU pRF parameters fits the HCP data and found that these NYU predictions explained above 70% of variance in the HCP BOLD signal. Third, we compared the two datasets in terms of polar angle meridian asymmetries in V1 cortical magnification. The NYU data showed the same pattern previously reported for the HCP data (Benson et al., 2021): more cortical surface area was dedicated to the horizontal than the vertical visual field meridian, and to the lower than the upper vertical visual field meridian.

### 4.1 Benefits of the NYU Retinotopy Dataset and Data Availability

The NYU Retinotopy Dataset described and visualized here consists of a large set of publicly available retinotopy data and can serve as a complement to the HCP 7T Retinotopy Dataset. All of the HCP data were acquired and processed in the same manner, thus any potential limits or biases in one participant’s data may apply to all of them. The same is true for the NYU data. For this reason, no matter how large one dataset is, it is important to have multiple, independent datasets to validate findings.

The NYU Retinotopy Dataset is publicly available for download (*to be made available with publication*) and contains comprehensive data for all 44 participants. The dataset includes anatomical images, raw and preprocessed functional images on the *fsnative* and *fsaverage* surfaces, *vistasoft* pRF model outputs on *fsnative* and *fsaverage* surfaces, and hand drawn V1, V2, V3, and hV4 ROIs for individual participants. Further, the organization and structure of the dataset adheres to BIDS (Brain Imaging Data Structure ((K. J. Gorgolewski et al., 2016). This means that the dataset is accessible, follows a clear and intuitive structure, and can be readily reanalyzed by others using scripts in which the inputs adhere to BIDS conventions. For example, it should be relatively simple to reanalyze the NYU Retinotopy Dataset with a different version of *fMRIPrep* or implement a different pRF model.

Next, we provide hand-drawn ROI labels for V1, V2, V3, and hV4 for each participant, facilitating further analyses of these areas by other researchers. We do not provide explicit ROIs for other regions, instead we provide *Jupyter Notebook* code and instructions for users to hand-define additional ROIs in this or other datasets using *Neuropythy*. The data were acquired for the whole brain, enabling investigation of maps in a variety of findings outside occipital cortex, such as the recent studies of retinotopy in hippocampus (Silson et al., 2021), thalamus (Arcaro et al., 2015), cerebellum (van Es et al., 2019), and parietal and frontal cortices (Mackey et al., 2017). Individual participant maps in the NYU Retinotopy Dataset extend well beyond hV4 into higher-order field maps. There are a number of competing hypotheses regarding the organization of higher-order visual areas that remain unresolved. The NYU Retinotopy dataset can be used as an additional dataset to help address competing hypotheses. For example these data may enable neuroscientists to resolve disagreements in regards to the retinotopic organization of maps in lateral occipital cortex (Amano et al., 2009; Kolster et al., 2010; Larsson & Heeger, 2006), and anterior ventral maps (Arcaro et al., 2009; Hansen et al., 2007; McKeefry & Zeki, 1997; Sereno et al., 1995; Wade et al., 2002; Winawer & Witthoft, 2015). Currently, two widely used atlases, Wang et. al. (2015) and Glasser et. al. (2016), have quite different parcellations of lateral and ventral occipital areas of visual cortex. Finally, the data could also be used to address questions of pRF shape, such as center surround organization (Zuiderbaan et al., 2012), ellipticity (Silson et al., 2018), or possibly validate recently identified areas in visual cortex (Elshout et al., 2018; Mikellidou et al., 2017).

As we have implemented the *vistasoft* pRF model on the HCP data, we also provide a new release of the *vistasoft* pRF model solutions for all 181 HCP participants on the *fsaverage* surface (*to be released with publication*). This complements the published pRF solutions from the same data using different software.

### 4.2 Vertex-wise polar angle, eccentricity, and pRF size estimates are well-correlated between datasets

First, we tested the vertex-wise similarity of NYU and HCP polar angle, eccentricity, and pRF size estimates using the parameter-median data. Overall, the pRF estimates were strongly correlated between the two datasets.

Vertex-wise polar angle estimates were almost perfectly correlated between the two datasets. Both the NYU and HCP data produced similar polar angle estimates at each vertex, and this was consistent across visual areas. Previously, a within-session split-half analysis of the HCP 7T Retinotopy Dataset identified polar angle estimates as the most internally reliable pRF parameter (Benson et al., 2018). Other work has shown that the polar angle estimates are consistent for the same participants across sessions and across aperture types (van Dijk et al., 2016). Here, we extend these analyses by showing that at the group level, the polar angle maps are highly similar between datasets.

Similarly, vertex-wise eccentricity estimates were strongly correlated, however, the two sets of estimates differed by a scale factor (~.75 in V1 to V3 and ~0.4 in hV4), with the NYU eccentricity estimates higher than those derived from the HCP data.

A scale factor difference in eccentricity between the datasets could arise for multiple reasons. First, it could be due to an anatomical shift of one dataset relative to the other. Specifically, one could account for the observed 3:4 ratio in eccentricity estimates by a translation along the posterior-anterior axis of about 4 mm, assuming the cortical magnification function of (Horton & Hoyt, 1991). Differences in the registration methods used during preprocessing might result in such a small, but systematic, shift in alignment: A cortical curvature-based alignment is used for the NYU data whereas a multimodal “*MSMAll*” alignment is used for the HCP data (Glasser et al., 2016). The *MSMAll* alignment includes a de-drifting relative to the *MSMSulc* algorithm, making a simple large-scale translation unlikely. Nonetheless, the differences in methods are likely to produce some differences in alignment. A second possibility is calibration errors in the stimulus size. In each dataset, it was assumed that all participants viewed the stimulus from a fixed distance (83.5 cm for NYU experiment and 101.5 cm for HCP experiment). Inevitably, there are small variations in the viewing distance, and thus stimulus size, due to differences in the positioning of each participant in the scanner. This has previously been noted in the methods of the HCP retinotopy study (Benson et al., 2018). Errors in stimulus size estimation would result in a scale factor difference in the eccentricity but not polar angle data, as is observed in our data. Third, differences in eccentricity estimates might be linked to differences in the pRF stimuli. The stimuli differed in temporal frequency (3 vs 15 Hz stimulus image update), width (3.1° vs 2°), and maximal extent (12.4° vs 8° radius). Differences in estimated eccentricity possibly arise from non-linearities in the neural response that are not modeled in the 2-D Gaussian pRF model. Finally, any of these three factors might combine to give rise to the observed differences.

Like eccentricity, pRF sizes were well-correlated between the two datasets, with a systematic bias towards larger pRFs in the NYU data. This bias was found in V1, V2, V3, and hV4. The simplest explanation is that the pRF size difference is caused by the same factors that cause the eccentricity difference. Supporting this, when pRF size is compared to eccentricity within a dataset, the differences between datasets are greatly reduced. For example, in both datasets, at 4° eccentricity, the pRF sizes are about 1° in V1 and are about 2° in V3. Nonetheless, some differences remain. In particular, the NYU pRF sizes were larger than the HCP sizes at central eccentricities (below 3°) but the pRF sizes were relatively similar in both datasets at more peripheral eccentricities (beyond 3° eccentricity). In accordance with a previous report (Linhardt et al., 2021), differences in pRF size may arise from the rotating wedge and expanding ring stimuli used in the HCP experiments, but not the NYU experiments. This is because the wedge and ring stimuli scale in size with eccentricity and may be useful for estimating small pRF sizes near the fovea.

In a supplemental analysis, we tested whether restricting the HCP data to bar scans alone would result in larger foveal pRF sizes. In V1 - hV4, HCP bar only pRF sizes were larger around foveal eccentricities when compared to HCP pRF sizes computed from the bar, wedge, and ring stimuli. We then tested whether using HCP bar only data could account for differences in pRF size between the NYU and HCP datasets. We found that V1 and V2 foveal HCP pRF sizes computed from HCP bar only data were *larger* than the NYU pRF sizes. The HCP pRF sizes in V3 were similar (although marginally smaller) than those in the NYU data, and this marginal difference was consistent across eccentricity. The HCP pRF sizes remained smaller in hV4 when compared to the NYU data, and this was also consistent at all eccentricities. These supplementary analyses show that using a bar stimulus *increases* pRF sizes around the fovea relative to using wedges, rings, and bars, and this suggests that there must be additional contributors (beyond the inclusion of the rotating wedge and expanding/contract ring stimuli) to the differences between NYU and HCP pRF sizes.

### 4.3 Differences between the NYU and HCP datasets in hV4

The hV4 map properties differ more between the two datasets than the V1, V2, and V3 map properties. Several factors may contribute to this difference. First, because the two datasets were aligned to the template surface using different algorithms, there is likely to be some degree of systematic spatial warping between the two datasets. A spatial warping will cause a relatively larger misalignment between small visual areas than large visual areas. This could explain the shallow slope in the hV4 vertex-wise scatter plots of both eccentricity and pRF size. Second, we compared map properties between the two datasets using visual field map ROIs defined by an anatomical template, rather than ROIs identified in individual participants based on functional data. Anatomical atlases tend to be most accurate for early visual areas, especially V1 (Benson et al., 2012), but also V2 and V3 (Benson et al., 2014; Benson & Winawer, 2018). They tend to be less accurate for higher visual areas such as hV4 (Wang et al., 2015). As a result, the hV4 comparisons are likely to include more data from other neighboring areas than the V1, V2, and V3 comparisons. Third, the two datasets differ in voxel size. The NYU voxels at acquisition are about two times greater in volume than the HCP voxels. Greater volume could result in larger pRF sizes, as PRF size depends on both neural receptive field size and the scatter of neural receptive field centers across the visual field (Dumoulin & Wandell, 2008). At acquisition, larger voxels contain more neurons and hence more scatter. Neural receptive field scatter will likely have a large effect on estimated pRF size for voxels in small visual areas. If a visual area is small (such as hV4, which is about half the size of the V1 to V3 maps) then the neurons within a single voxel will sample a relatively large portion of the visual field, hence there will be more scatter per voxel. The effect of voxel size on pRF size, however, is potentially complicated by pre-processing procedures, especially the spatial resampling of the fMRI data to the surface. For example, the HCP pRF analyses were conducted on a spatially downsampled mesh, which could blur the time series and affect pRF size.

### 4.4 Fitting predicted time-series based on the NYU pRF parameter estimates to the HCP BOLD percent signal change

We assessed cross-dataset reproducibility by comparing vertex-wise pRF parameters between the NYU and HCP datasets. A good model should also be able to predict the time-series data, in addition to the estimated parameters. If the estimated parameters were identical, then it would trivially follow that the predicted time-series would also match. However, the relation between pRF parameters and the time-series is non-linear. Thus, the similarity of the time-series between the datasets is not directly inferred from the similarity of the pRF parameters. We showed that indeed, the NYU pRF dataset could be used to predict the HCP BOLD signal of the vertices within the V1-hV4 maps, with high variance explained (more than 70%). This is a strong test of generalization, as there are many differences between the datasets, such as wedge and ring stimuli in the HPC data, and only bar stimuli in the NYU data.

Finally, the variance explained metric provides a condensed overview of the cross-dataset reproducibility. This metric takes three parameter comparisons (polar angle, eccentricity, and size) and reduces it to one (variance explained, *R*^2^). This single metric is easily visualized as an overview map, which shows high variance explained in visual cortex, especially V1-hV4, but also nearby lateral, temporal, and dorsal regions, suggesting that cross-dataset reproducibility extends outside the regions tested here.

### 4.5 Implications of the similarities and differences between the NYU and HCP Datasets

The pattern of similarities and differences between the two datasets is important for quantitative interpretation of retinotopic data, especially with respect to comparisons among data from different sites. The polar angle data are highly consistent between the NYU and HCP experiments, with circular correlation coefficients above 95% in each of V1, V2, V3, and hV4. Polar angle is especially important for identification of visual area boundaries, thus different studies are likely to be able to achieve close agreement in delineation of visual field maps. Why do the polar angle estimates show such close agreement? First, polar angle estimates are immune to differences in scale of stimulus coordinates. Second, they are likely to be insensitive to differences in voxel size. Third, pRF model predictions, given typical stimulation sequences, are sharply tuned to polar angle, enabling precise estimates of this parameter (Lage-Castellanos et al., 2020; Lerma-Usabiaga et al., 2020).

The small but systematic differences in eccentricity and pRF size indicate that a measure of tolerance is required when generalizing from one dataset to another. For example, the anatomically defined retinotopic template from Benson and Winawer (2018) was generated using the HCP data. If the template is applied to a different dataset without constraints from functional measures, there might be a small-scale factor error in the eccentricity estimates in V1, V2, and V3, and a larger error in hV4 and other higher visual areas. The fact that the parameter estimates are so strongly correlated between the datasets indicates that random noise plays a minimal role in generating the differences in eccentricity and pRF size seen here. In turn, this suggests that it should be feasible to resolve the modest between-dataset differences, for example, with a corrective spatial warping or a calibration using a standardized stimulus and anatomical landmarks. However, we do not currently know where ground truth lies relative to either dataset, thus it is not prudent to warp one dataset to match the other.

### 4.6 What is ground truth?

Discrepancies between the NYU and HCP data should not be taken as proof that either of the datasets is the more precise one. There are multiple possible reasons for discrepancies in pRF estimates when comparing between two datasets that differ in so many ways.

First, a pRF is a population measure. If the underlying neural populations differ, their receptive fields should differ as well. For example, a larger voxel is expected to have a larger pRF; this is not a measurement error, but rather an inherent property of population measures. Likewise, larger voxels are unlikely to have pRF centers at extreme values, such as those along vertical meridian, because any pooling will tend to push the measure toward the population average. This too is not a measurement error, but rather a property of population measures. Second, pRF models (like all models) are approximations of more complex underlying systems. The pRF models used here, and in the HCP paper (Benson et al., 2018), make predictions based on the stimulus apertures, and do not account for either the spatial or temporal properties of the stimulus carrier, nor the task the participant performs, both of which are known to influence the neural responses (Le et al., 2017; Wandell & Winawer, 2015). Whereas retinotopic maps are not very flexible (Liu et al., 2006), pRF parameters do depend upon a variety of factors that are frequently not modeled (Fang et al., 2008). Even at the level of individual V1 neurons, when a linear receptive field is fit to neural data, the shape differs depending on the mapping stimuli (Victor et al., 2006). Thus, it is reasonable to expect experimental differences to result in differences in pRF parameters. By quantifying pRF similarity across two retinotopy datasets that were acquired using different methods, our work provides a reasonable estimate of how similar pRF parameters are likely to be when experiments differ across many dimensions.

Finally, we note that we do not have access to ground truth (i.e. the underlying true retinotopic arrangement in each participant’s visual cortex). Some systematic differences in pRF results are expected due to the nature of population measures and the simplifying assumptions of the models. Other differences may arise from preprocessing differences (e.g., registration methods). In principle, it may be possible to assess which preprocessing method better represents the ground truth data, however, this is only possible given access to ground truth data. This may be possible through simulation methods such as those used to validate pRF software (Lerma-Usabiaga et al., 2020), but such methods do not yet exist for assessing retinotopic maps themselves.

### 4.7 Polar angle meridian asymmetries in V1 cortical magnification are reproducible across datasets

Reports of *performance fields* show that visual performance on a range of tasks changes as a function of polar angle (Abrams et al., 2012; Baldwin et al., 2012; Barbot et al., 2021; Cameron et al., 2002; Carrasco et al., 2001; Carrasco et al., 2002; Corbett & Carrasco, 2011; Fortenbaugh et al., 2015; Fuller et al., 2008; Greenwood et al., 2017; Himmelberg et al., 2020; Kurzawski et al., 2021; Levine & McAnany, 2005; Lundh et al., 1983; Montaser-Kouhsari & Carrasco, 2009; Pointer & Hess, 1989; Regan & Beverley, 1983; Rijsdijk et al., 1980; Robson & Graham, 1981; Silva et al., 2008; Talgar & Carrasco, 2002; Virsu & Rovamo, 1979). Previous fMRI studies have reported similar polar angle asymmetries in V1 BOLD amplitude (Liu et al., 2006; O’Connell et al., 2016), cortical magnification (Benson et al., 2021; Silva et al., 2018), and pRF sizes (Silva et al., 2018). In parallel to behavior, more cortical space is dedicated to processing the horizontal than vertical visual field meridian (i.e., a cortical HVA). Likewise, there is more cortical space dedicated to the lower vertical than upper vertical visual field meridian (i.e., a cortical VMA). We tested whether these polar angle asymmetries in V1 surface area are also found in the NYU data.

The HVA was identified in the NYU data; around a 65% HVA asymmetry index for a ±15° wedge. Thus, there is almost twice as much cortical space along the horizontal than the vertical visual field meridian. The HVA became progressively weaker with increasing wedge-width. Thus, the cortical HVA is strongest along the meridians themselves. This measurement qualitatively and quantitatively corresponds to the cortical HVA identified by Benson et al. (2021) in the HCP data. Likewise, the VMA was identified in the NYU data; around a 30% VMA asymmetry index for a ±15° wedge. Thus, there is more surface area dedicated to the lower than upper visual field meridian. This is similar to the cortical VMA identified by Benson et al. (2021), however, one discrepancy is that in the NYU data, the VMA asymmetry decreased with wedge-width more gradually than in the HCP data.

Notably, our cortical HVA and VMA measurements are commensurate with recent psychophysical data, both of which show a larger HVA than VMA. For contrast sensitivity, the HVA is approximately 60% (similar to the ~65% cortical HVA reported here) whereas the VMA is weaker at approximately 20% (similar to the ~30% cortical HVA reported here) (Himmelberg et al., 2020). For acuity, the HVA is approximately 40% and the VMA is approximately 20% (Barbot et al., 2021). Our cortical magnification measurements match these perceptual measurements; in fact, the contrast sensitivity measurements from Himmelberg et al. (2020) come from a subsample of the participants included in the present study. Recent implementations of computational models have shown that polar angle asymmetries in photoreceptor and retinal ganglion cell sampling density only account for a small portion of *performance field* asymmetries, and that the sampling of the visual field as a function of polar angle is more asymmetric in cortex than in the retina (Kupers et al., 2019, 2020). We are currently developing a computational model to assess the extent to which these cortical asymmetries can account for perceptual performance field asymmetries in contrast sensitivity.

Overall, the HVA is strong and clear, and these asymmetries in cortical geometry can be measured robustly in independent datasets. Qualitatively, the VMA was also reproduced between the two datasets, corroborating that this cortical asymmetry exists. However, there were small differences in the detailed characterization of the VMA between the two datasets. The asymmetry between the upper and lower meridian declined faster with angular distance from the vertical meridian in the HCP dataset than in the NYU dataset. Such differences may arise from subtle differences in our analysis compared to Benson et al. (2021). In this study, we examined the surface area within V1 only, whereas Benson et al. (2021) accounted for vertical meridian surface area in both V1 and V2. Additionally, the methods used to define the wedge, and the sizes of the wedges themselves, differed between the two studies. Further, the NYU data used cleaned polar angle data (see *2.9.2 Eccentricity boundaries for sub-ROIs*), whereas Benson et al. (2021) used raw polar angle data. Thus, it is not surprising that small differences in results arise from small differences in analysis. Nonetheless, despite some quantitative differences, the overall pattern of more surface area for the horizontal than the vertical visual field meridian representation, and for the lower than the upper vertical visual field meridian representation, holds across datasets and analysis methods.

### 4.8 Retinotopic maps are reproducible

Identifying and understanding retinotopic maps is a central aim of visual neuroscience. We endeavor to understand how the visual brain is organized into multiple visual areas, identify the functional roles of these areas, and understand how they unite to make sense of the influx of sensory information that ultimately gives rise to visual perception. This is no small feat, and if newly identified retinotopic areas and features are to be considered generalizable across the human population, they must be subject to tests of validity and reliability to ensure that findings are robust and generalizable across independent datasets.

Previous studies have assessed reproducibility of retinotopic maps via different tests of validity. PRF estimates are reported to have good intersession reliability. Specifically, they are consistent across time in the same observers who were measured twice in the same scanner (Lage-Castellanos et al., 2020; van Dijk et al., 2016). This is supported by split-half analysis in the HCP retinotopy data, in which each participant’s functional scans were split into first and second halves, and pRF parameters were estimated twice (Benson et al., 2018). Likewise, pRF estimates have been shown to be similar across participants when measured with the same stimuli, in the same scanner, and analyzed from the same data; the HCP data analyzed as split half by participants showed that each subgroup of half the participants had similar average pRF maps (Benson et al., 2018). Furthermore, it has been shown that pRFs are reproducible when the same participants are tested with different stimulus apertures and carrier images, using the same scanner and same software (van Dijk et al., 2016). Finally, computational validity is the ability of software to reproduce the *correct result* for simulated ground truth test data (irrespective of whether it is the *same* answer), and this has been assessed using a recent validation framework tested against different pRF modelling software (Lerma-Usabiaga et al., 2020). These studies have addressed the reproducibility of retinotopic maps *within* their own datasets. Here, we have assessed retinotopic reproducibility in a different way from these prior studies. We assessed reliability of retinotopic maps at the group level, by comparing two *independent* datasets that differed in their stimulus apertures, participants, MRI hardware, fMRI protocol, and preprocessing pipeline. Many experimental differences are typical for retinotopy data collected from different labs or sites, and up to now it was unknown how similar pRF estimates were when many elements of experimental design were simultaneously altered. Here, we have shown that fundamental retinotopic properties in human V1 to hV4 can be reliably reproduced in independent datasets that simultaneously differ in many aspects.

There is much considerable interest in the reproducibility of scientific results in the biomedical sciences (Ioannidis, 2005), with attention drawn to results in psychology (Open Science Collaboration, 2015) and neuroimaging (Poldrack et al., 2017). For example, the same data analyzed by different groups often led to different results and conclusions (Botvinik-Nezer et al., 2020). Similarly, reports show that reproducibility is poor for many task-based fMRI experiments (Elliott et al., 2020). However, this may reflect limitations in the kind of experiments tested, as some studies show higher reproducibility than others (Kragel et al., 2021), and differences in analysis, as surface-based cognitive neuroscience studies of cerebral cortex (such as the current study) perform well with regards to reproducibility (Assem et al., 2021).

Our data show that human retinotopic maps are reliable and reproducible between independent datasets that have been acquired using different fMRI protocols, measured using different stimuli, and have undergone different preprocessing. This is likely because the method underlying the measurement of retinotopic maps circumvents many of the issues involved in irreproducibility (see Poldrack et al. (2017) for review). PRF estimates are computed using an explicit computational model of the fMRI response and this model is defined in terms of input parameters (Dumoulin & Wandell, 2008; Wandell & Winawer, 2015). These input parameters are grounded in solid theory of neural visual receptive fields; a pRF is formalized as a 2D-Gaussian and this is based on the established physiological structure of neural receptive fields in the visual cortex (Hubel & Wiesel, 1968). Thus, the fMRI response is modelled using a quantitative characterization of neural activity. Using this powerful computational modeling approach, retinotopic maps can be reliably measured, and integrated, across different neural measurements (Wandell, 1995; Wandell et al., 2015).

## 5. Conclusion

We have tested the cross-dataset reproducibility of retinotopic properties by comparing two independent datasets that differ in many aspects. We have reported that polar angle, eccentricity, and pRF size estimates are well-correlated between the two datasets, albeit with systematically greater eccentricity and pRF size estimates in the NYU data. These systematic differences in eccentricity and pRF size might be linked to small differences in anatomical alignment methods or viewing distance. Differences in pRF size may also arise from differences in voxel size and stimulus aperture. Next, we identified polar angle asymmetries in V1 cortical magnification similar to those found in the HCP data. There was more cortical surface area dedicated to the horizontal than vertical visual field meridian, and to the lower than upper vertical visual field meridian, and these asymmetries decreased gradually with increasing angular distance from the meridians. The pattern and strength of these asymmetries was similar to those reported from previous behavioral measurements, supporting a link between brain and behavior. Overall, we have shown that we can reliably describe retinotopic properties in human V1 to hV4 from two datasets that differ in many ways, and we highlight the importance of the cross-dataset validation of new retinotopic findings to ensure that they can be generalized across the human population. The NYU Retinotopy Dataset can serve as a benchmark for testing hypotheses about the organization of human visual cortex and for comparison to the HCP 7T Retinotopy Dataset.

## Author contributions

M.M.H, J.W.K, M.C, and J.W conceived the experiments. M.M.H, N.C.B, and J.W performed the experiments. M.M.H, J.W.K, & J.W analyzed the data. M.M.H and J.W wrote the paper with feedback from J.W.K, N.C.B, D.G.P, and M.C.

## Acknowledgements

Some of the NYU datasets were originally collected for other projects for which retinotopy was just one component. We thank Billy Broderick, Serra Favila, Iris Groen, and Eline Kupers for contributing their retinotopy datasets to this paper. This research was funded by the US National Eye Institute R01-EY027401 to MC and JW and R01-EY027964 to DGP and JW.

## Supplementary Materials

**Figure S1.**
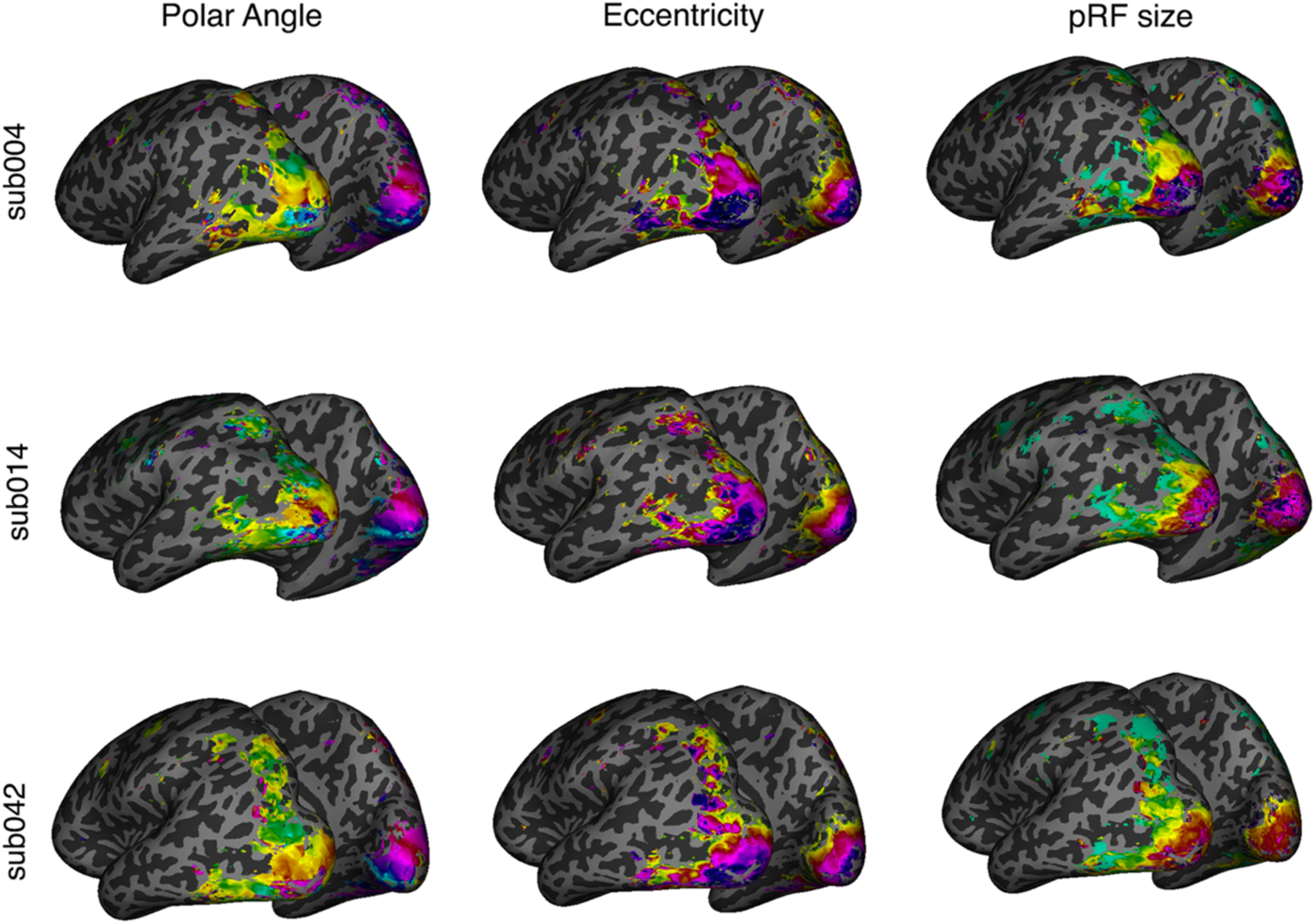
Examples of retinotopic maps for three individual participants (subj004, subj014, and subj042) projected onto their inflated *fsnative* surfaces.

**Figure S2.**
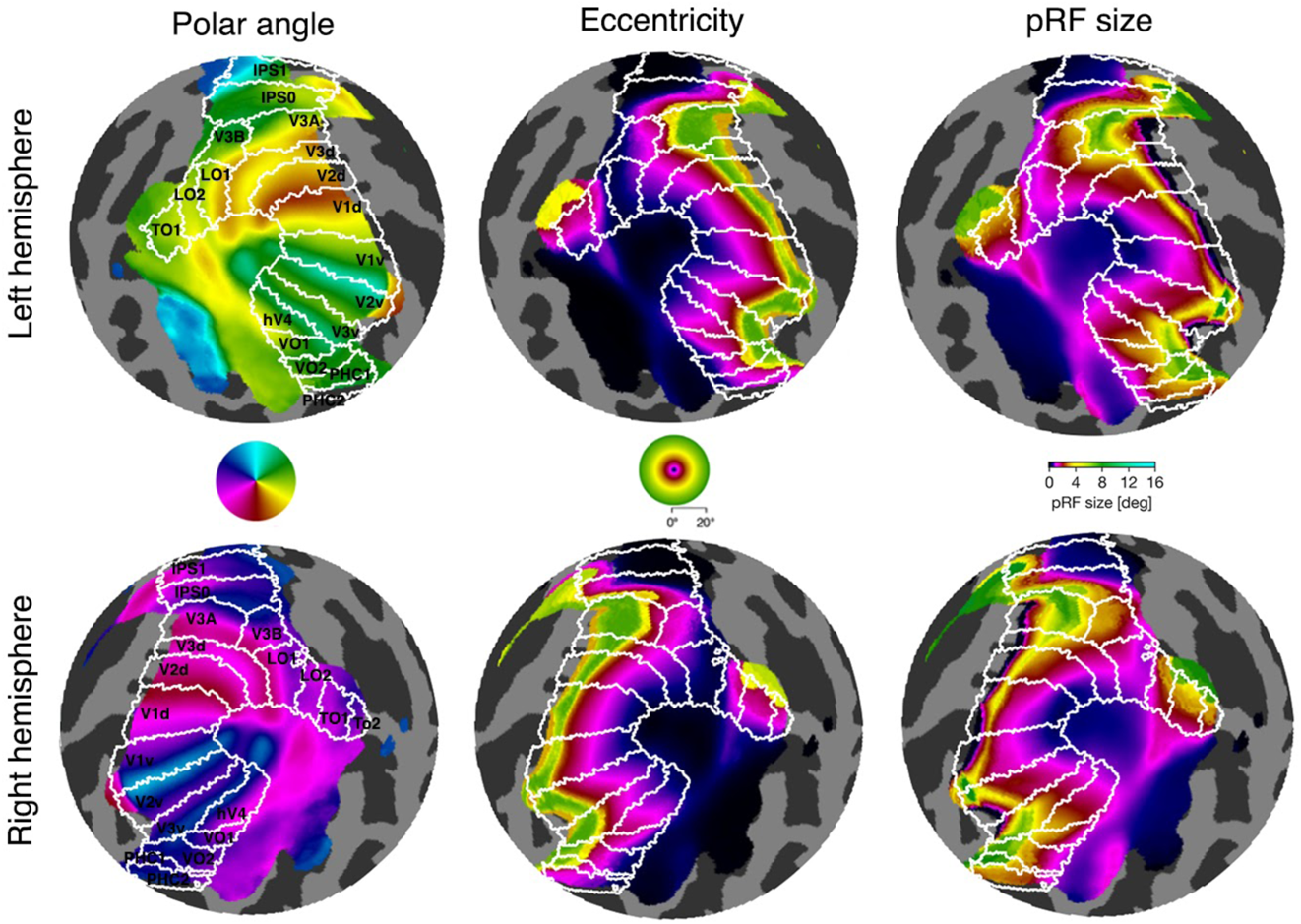
Retinotopic maps for the HCP group-average time-series data projected on *fsaverage* flatmaps for the left and right hemisphere. White boundaries specify ROIs derived from the Wang Maximum Probability Atlas.

**Figure S3.**
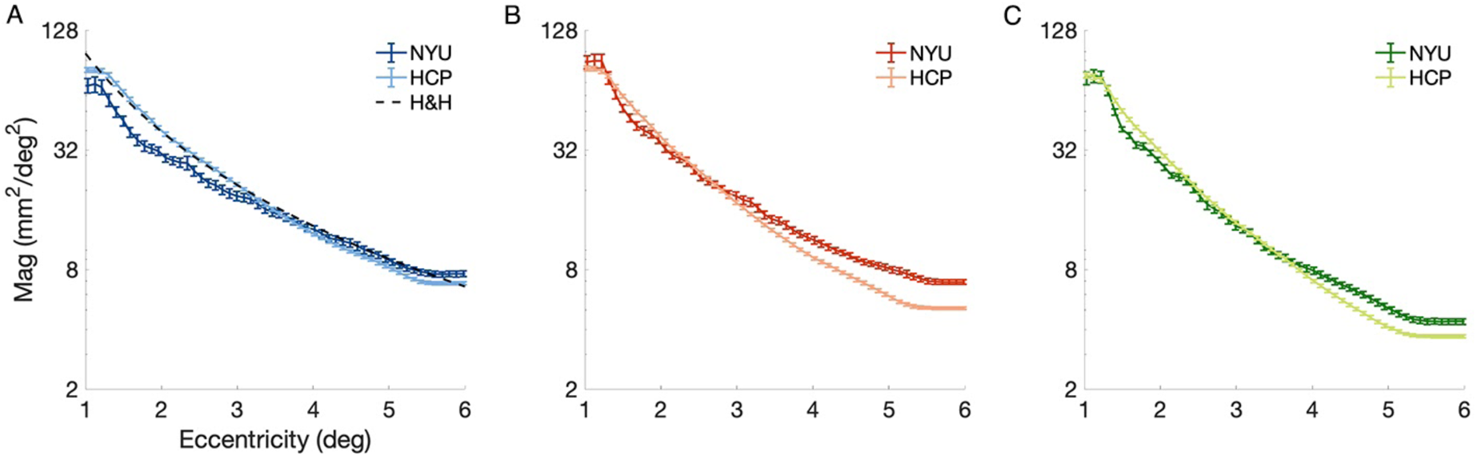
Areal cortical magnification plotted as a function of eccentricity, averaged across individual participants for the NYU (dark lines) and HCP (light lines) datasets, in V1, V2, and V3. Cortical magnification is calculated on the *fsnative* surface for each participant. The black dashed line in V1 represents the cortical magnification function for V1 as reported by Horton and Hoyt (1991). Error bars are ±1 SEM.

**Figure S4.**
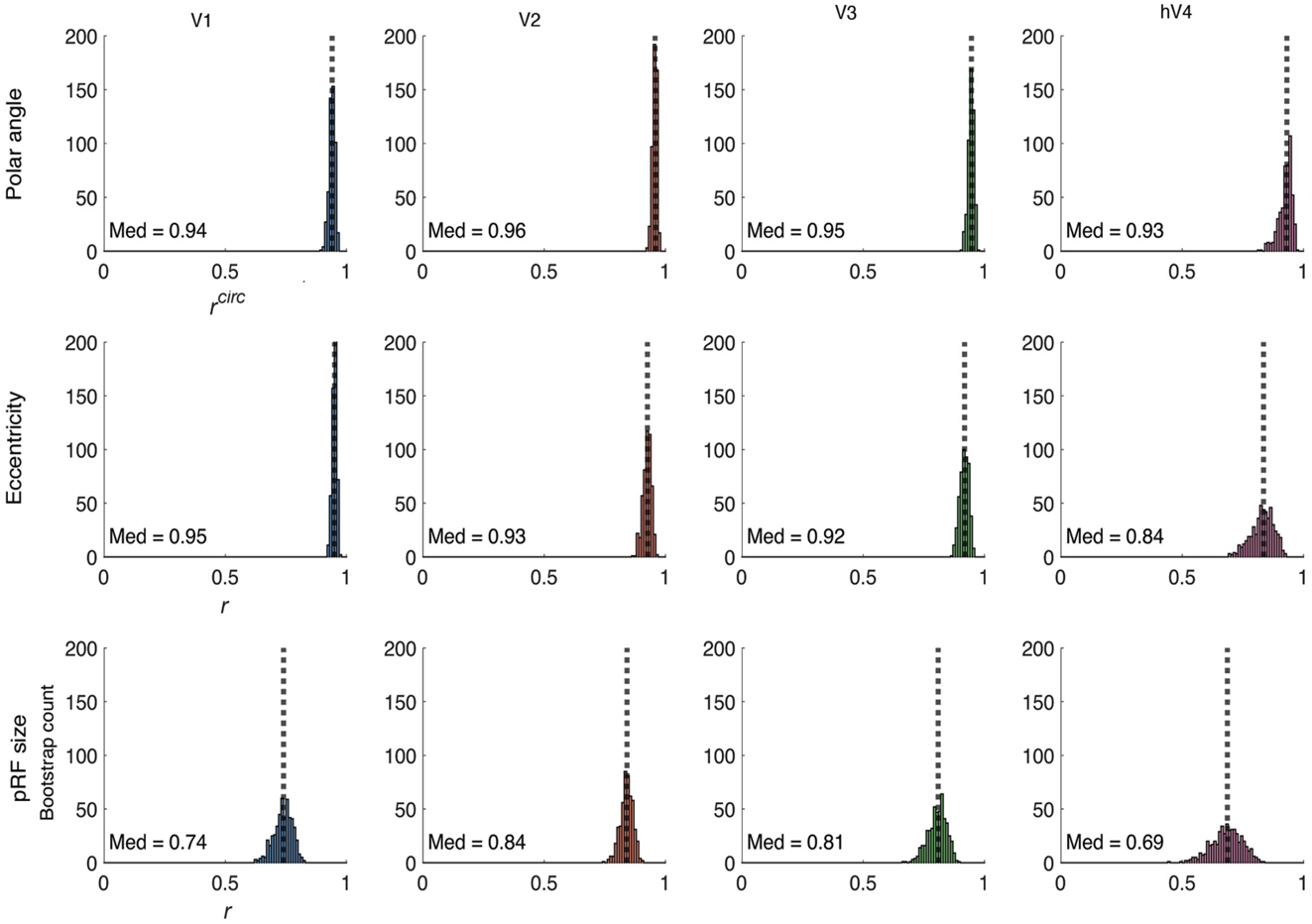
Histograms of 500 subsampled correlations for V1 to hV4, for polar angle, eccentricity, and pRF size estimates. Each iteration draws 20 participants from the NYU and 20 participants from the HCP dataset (with replacement), computes median-parameter maps for the subsampled NYU and HCP data, yielding a single *r* value per iteration for each parameter (polar angle, eccentricity, size) and for each visual field map. The dotted lines represent median *r* value across the 500 iterations.

**Figure S5.**
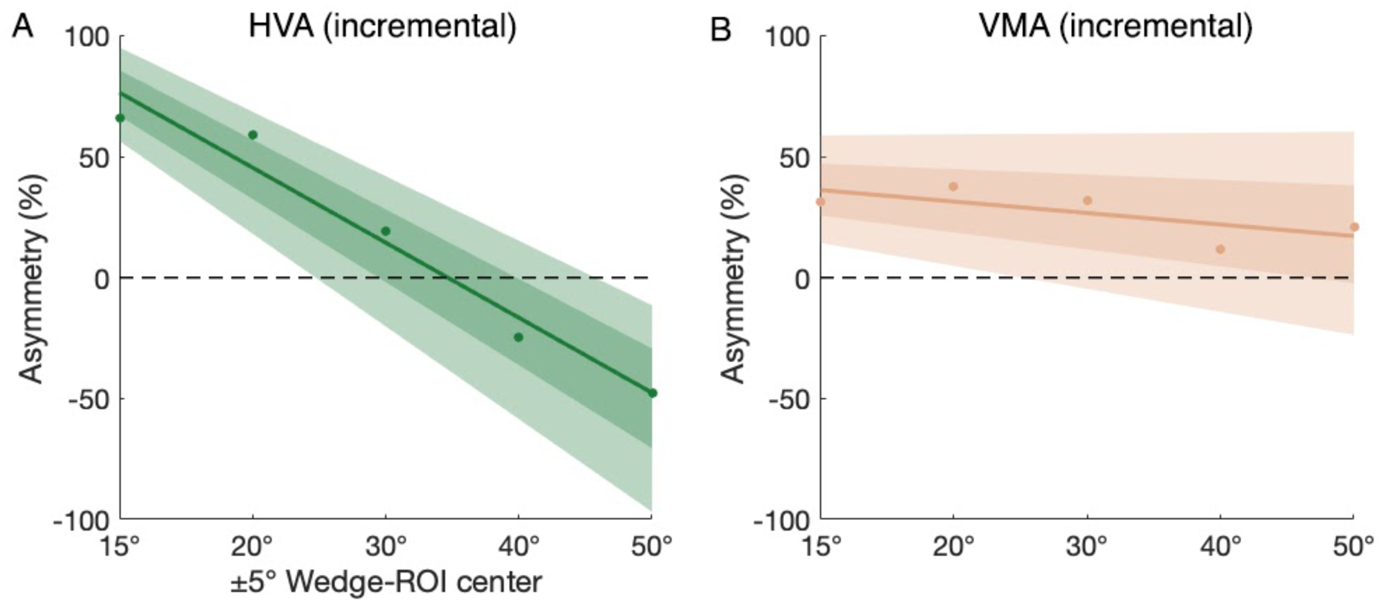
Incremental HVA and VMA asymmetries for a ±5° wedge centered at 5 different locations from the visual field meridians. The HVA and VMA decrease with increasing distance from a cardinal meridian. The incremental HVA becomes inverted because a ±5° wedge centered 50° from the horizontal meridian is closer to the vertical than the horizontal meridian (i.e., beyond 45°). This is the same for a ±5° wedge centered 50° from the vertical meridian; It will be closer to the horizontal than the vertical meridian, so the flip is expected.

**Figure S6.**
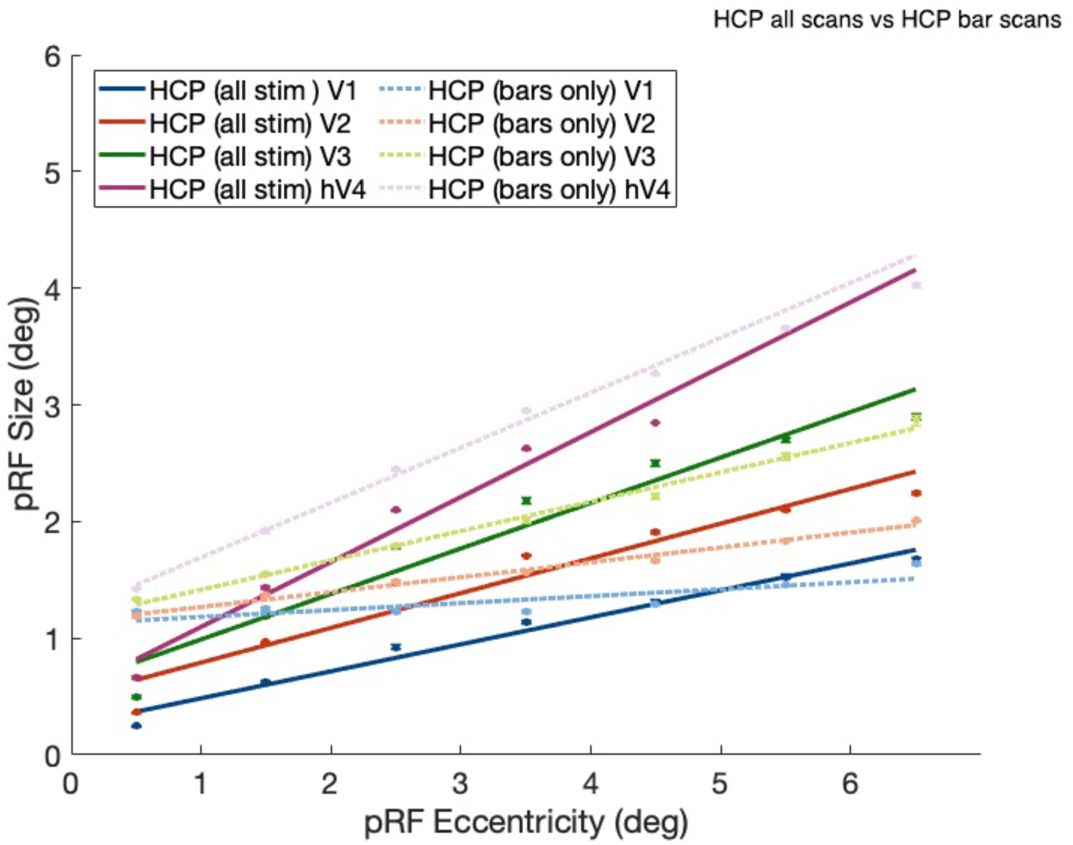
Parameter-median PRF sizes as a function of eccentricity for the HCP data using bars, wedges, and rings (solid lines) and the HCP data using bars alone (dashed lines) for V1, V2, V3, and hV4. pRF sizes are binned into 1° eccentricity bins and a linear function is fit to the mean sizes. Error bars are ±1 SEM (vertices per bin), R^2^ threshold >= 0.4.

**Figure S7.**
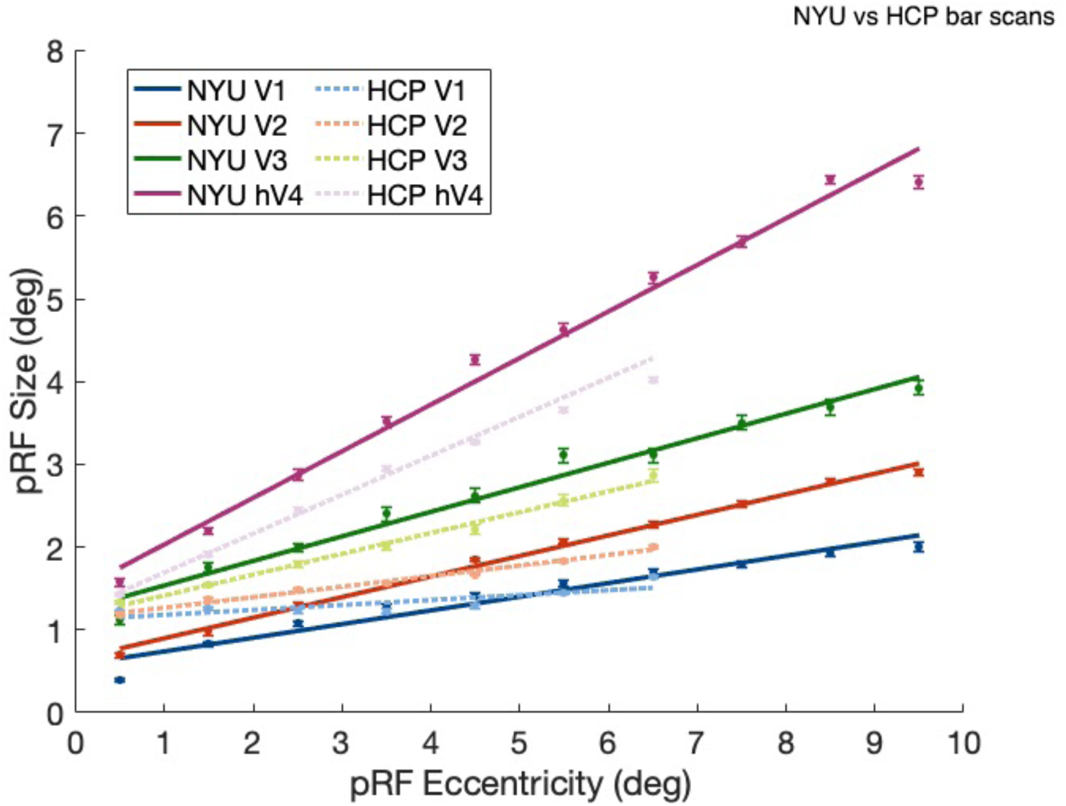
Parameter-median PRF sizes as a function of eccentricity for the NYU data and the HCP data using bars alone (dashed lines) for V1, V2, V3, and hV4. pRF sizes are binned into 1° eccentricity bins and a linear function is fit to the mean sizes. Error bars are ±1 SEM (vertices per bin), R^2^ threshold >= 0.4.

**Figure S8.**
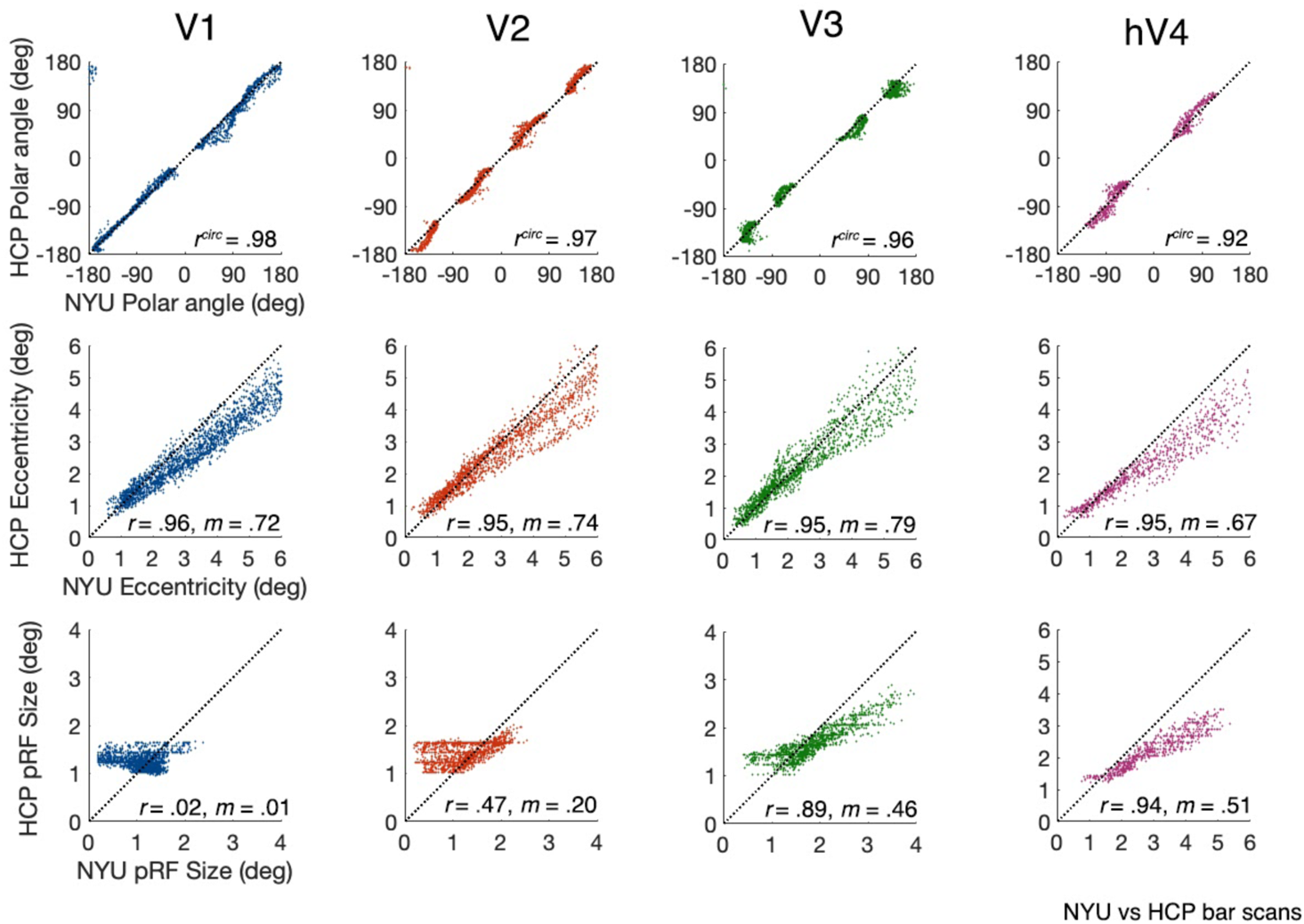
Scatterplots comparing vertex-wise parameter-median pRF estimates from the NYU data and the HCP bar only data within V1, V2, V3, and hV4. Each datapoint represents an *fsaverage* vertex. The black dashed line represents *y* = *x*. All reported *r* values are highly significant (*p* < .001) except for V1 pRF size (*p* > .05). *m* represents slope.

**Figure S9.**
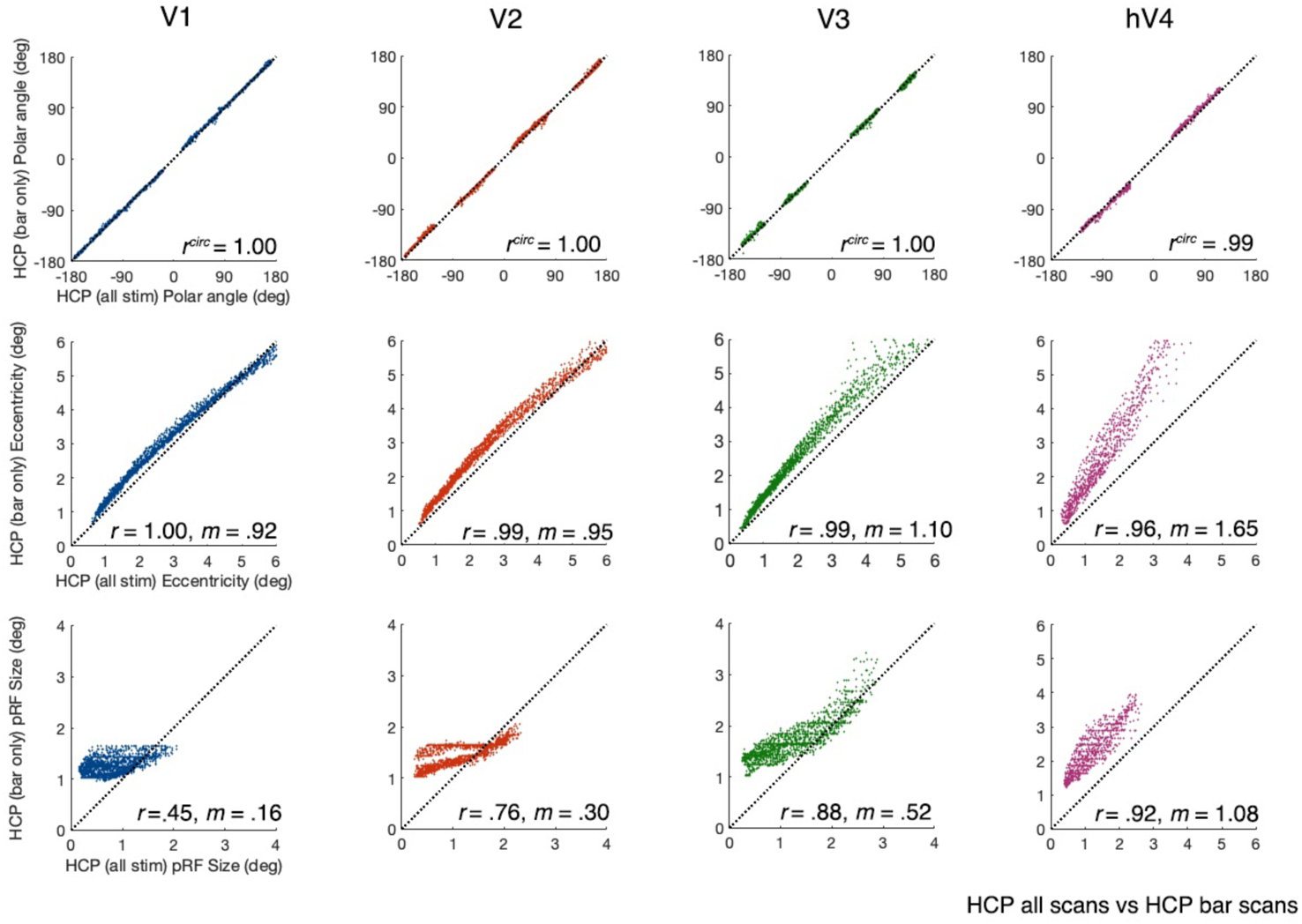
Scatterplots comparing the vertex-wise parameter-median pRF estimates from the HCP bars, wedges, and rings data, and the HCP bar only data, within V1, V2, V3, and hV4. Each datapoint represents an *fsaverage* vertex. The black dashed line represents *y* = *x*. All reported *r* values are highly significant (*p* < .001). *m* represents slope.

## Notes

**Conflict of interest:** The authors declare no competing financial interests.

https://osf.io/e6vqk/

### Competing Interest Statement

The authors have declared no competing interest.

### Summary of Updates

new analyses text revisions

